# Early microglial response, myelin deterioration and lethality in mice deficient for very long chain ceramide synthesis in oligodendrocytes

**DOI:** 10.1101/2022.05.29.493337

**Authors:** Jonathan D Teo, Oana C Marian, Alanna G Spiteri, Madeline Nicholson, Huitong Song, Jasmine XY Khor, Holly P McEwen, Laura Piccio, Jessica L Fletcher, Nicholas JC King, Simon S Murray, Jens C Brüning, Anthony S Don

## Abstract

The sphingolipids galactosylceramide (GalCer), sulfatide (ST) and sphingomyelin (SM) are essential for myelin stability and function. GalCer and ST are synthesized mostly from C22-C24 ceramides, generated by Ceramide Synthase 2 (CerS2). To clarify the requirement for C22-C24 sphingolipid synthesis in myelin lipid biosynthesis and stability, we generated mice lacking CerS2 specifically in myelinating cells (CerS2^ΔO/ΔO^). At 6 weeks of age, normal-appearing myelin had formed in CerS2^ΔO/ΔO^ mice, however there was a reduction in myelin thickness and the percentage of myelinated axons. Pronounced loss of C22-C24 sphingolipids in myelin of CerS2^ΔO/ΔO^ mice was compensated by greatly increased levels of C18 sphingolipids. A distinct microglial population expressing high levels of activation and phagocytic markers such as CD64, CD11c, MHC class II, and CD68 was apparent at 6 weeks of age in CerS2^ΔO/ΔO^ mice, and had increased by 10 weeks. Increased staining for denatured myelin basic protein was also apparent in 6-week-old CerS2^ΔO/ΔO^ mice. By 16 weeks, CerS2^ΔO/ΔO^ mice showed pronounced myelin atrophy, motor deficits, and axon beading, a hallmark of axon stress. 90% of CerS2^ΔO/ΔO^ mice died between 16 and 26 weeks of age. This study highlights the importance of sphingolipid acyl chain length for the structural integrity of myelin, demonstrating how a modest reduction in lipid chain length causes exposure of a denatured myelin protein epitope and expansion of phagocytic microglia, followed by axon pathology, myelin degeneration, and motor deficits. Understanding the molecular trigger for microglial activation should aid the development of therapeutics for demyelinating and neurodegenerative diseases.

**Main Points:**

- Oligodendrocytes lacking CerS2 produce myelin using sphingolipids with C16/C18 instead of C22/C24 N-acyl chains
- C22/C24 myelin sphingolipids are essential for myelin stability, microglial quiescence, and survival beyond young adulthood

**Table of Contents Image:** 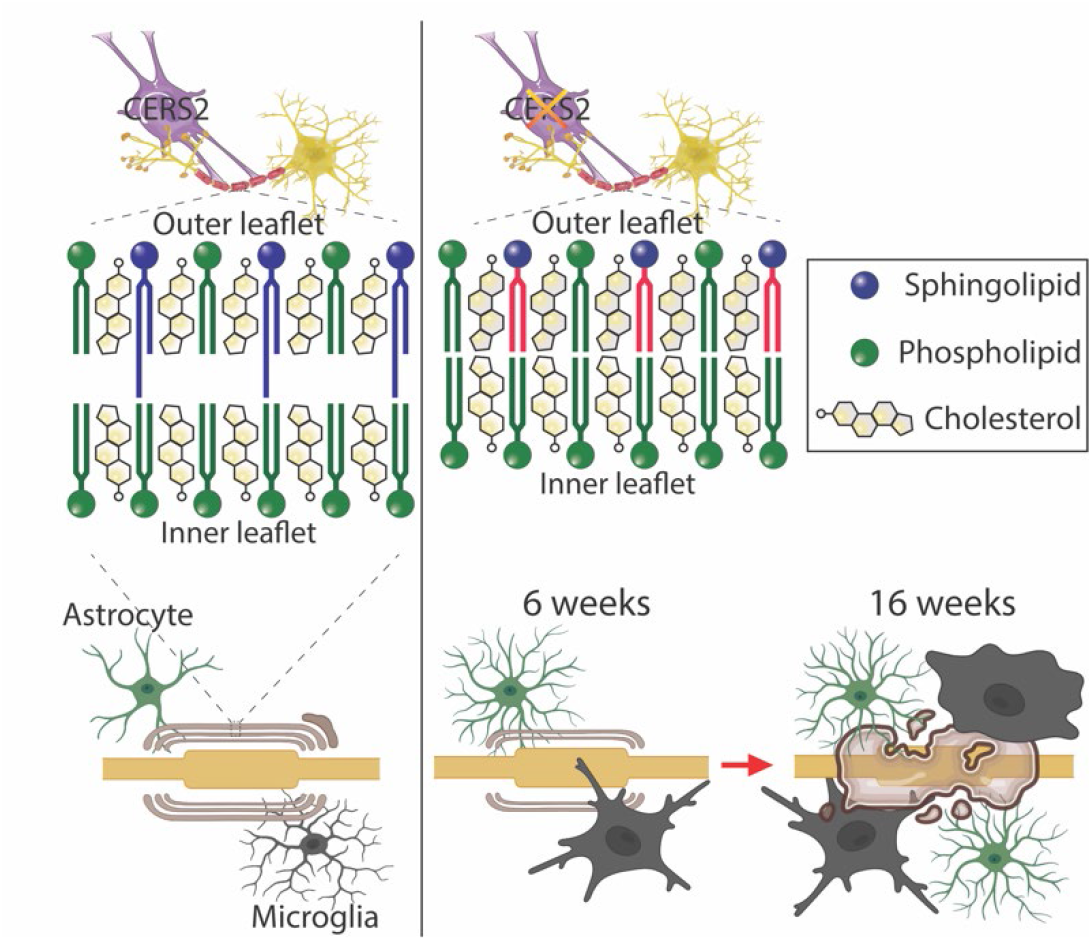

## Introduction

Myelin is a multi-layered extension of the oligodendrocyte or Schwann cell membrane that is wrapped in a spiral around neuronal axons (Simons & Nave, 2016). Myelin restricts the passage of ions across axonal membranes to the nodes of Ranvier, facilitating the fast and energy-efficient form of neuronal signalling termed saltatory conduction, which is essential for movement, sensory perception, and cognition (Fields, 2008).

At 70-80% lipid by dry weight, myelin is a particularly lipid-rich biological membrane (O’Brien & Sampson, 1965; Schmitt et al., 2015). The sphingolipids sphingomyelin (SM), galactosylceramide (GalCer), and sulfatide (ST), which are all synthesized from ceramides, make up 25-35% of myelin lipid in humans (O’Brien & Sampson, 1965; Quarles et al., 2006). SM and GalCer are produced by transfer of a choline phosphate or galactose moiety, respectively, to the primary hydroxyl of ceramide. ST is produced by sulfation of GalCer (Schmitt et al., 2015) (Fig. 1). While SM is present in all cells, GalCer and ST are relatively unique to myelin and essential for its stability. Mice lacking GalCer and ST due to genetic ablation of ceramide galactosyltransferase exhibit myelin instability, conduction deficits, progressive hindlimb paralysis, and early lethality (Coetzee et al., 1996; Dupree et al., 1998).

**Figure 1.**
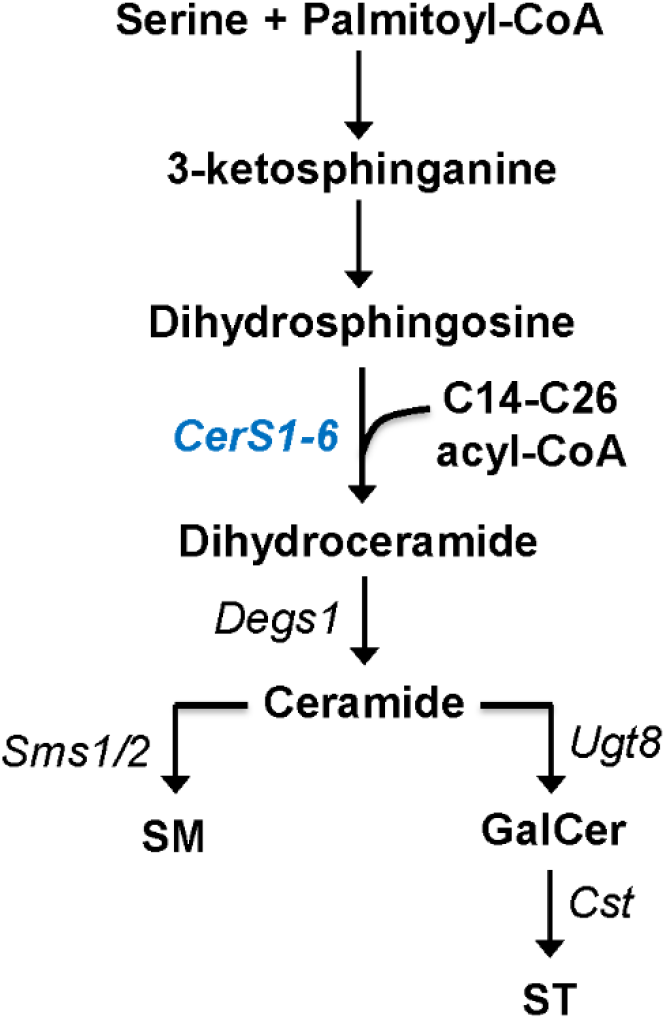
*De novo* sphingolipid synthesis in oligodendrocytes. Serine and palmitoyl-CoA are converted to 3-ketosphinganine, then dihydrosphingosine, through the actions of serine palmitoyl transferase and 3-ketosphinganine reductase. CerS1-6 use dihydrosphingosine and fatty acyl-CoA to produce dihydroceramides, which are desaturated to ceramides by sphingolipid Δ4-desaturase (Degs1). Ceramides can be converted into sphingomyelin (SM) by sphingomyelin synthases (Sms1/2), or galactosylceramide (GalCer) by UDP-galactose ceramide galactosyltransferase (Ugt8) and sulfatide (ST) by cerebroside transferase (Cst). These sphingolipids are crucial and abundant constituents of myelin. Alternatively, ceramide may be phosphorylated, deacylated by ceramidases to form sphingosine, or glucosylated by glucosylceramide synthase and further converted to gangliosides (Marian et al., 2020).

*De novo* ceramide synthesis in mammals proceeds through the transfer of a variable length fatty acid (usually 14-26 carbons; i.e. C14-C26) from fatty acyl coenzyme A (CoA) to the free amine of dihydrosphingosine, forming dihydroceramides (Fig. 1). This reaction is catalysed by a family of six ceramide synthases (CerS1-6), each of which exhibits a preference for particular fatty acyl CoA chain lengths (Park et al., 2014). Dihydroceramides are rapidly desaturated by sphingolipid Δ4-desaturase to form ceramides. GalCer and ST are preferentially synthesized from very long chain (C22-C24) ceramides produced by CerS2 in oligodendrocytes (Becker et al., 2008; Ben-David et al., 2011; Couttas et al., 2016). C18 ceramides, synthesized by CerS1, are also highly abundant in the brain and essential for ganglioside synthesis and neuronal viability during development (Ginkel et al., 2012).

Studies to date have not clarified whether the N-acyl chain length of myelin sphingolipids has a significant bearing on myelin stability. Mice bearing a gene trap disruption of the *Cers2* gene present with severe hepatomegaly, liver pathology and glucose dyshomeostasis (Park et al., 2013; Pewzner-Jung, Brenner, et al., 2010; Pewzner-Jung, Park, et al., 2010), as well as loss of myelin and cerebellar ataxia (Ben-David et al., 2011; Imgrund et al., 2009). These mice show almost complete loss of C22-C24 ceramides in liver and brain (Ben-David et al., 2011; Imgrund et al., 2009; Pewzner-Jung, Park, et al., 2010). This results in an overall reduction in hexosylceramide (HexCer) and ST content, associated with progressive myelin degeneration over several months (Ben-David et al., 2011; Imgrund et al., 2009). Median survival in CerS2-null mice was 5 months (Pewzner-Jung, Brenner, et al., 2010), however this phenotype could not be attributed to loss of CerS2 in any specific organ or physiological system due to the ubiquitous nature of the knockout. A case of progressive myoclonic epilepsy attributed to haploinsufficiency of the CerS2 gene was reported in 2014 (Mosbech et al., 2014), indicating that balanced levels of this enzyme activity are required for normal neurological function in humans.

Herein, we generated mice specifically lacking CerS2 in myelinating cells to determine how altered sphingolipid acyl chain composition, specifically within myelin, affects myelin stability and glial homeostasis in white matter. Deletion of CerS2 in oligodendrocytes caused a reduction in the acyl chain length of myelin sphingolipids without an overall reduction in myelin sphingolipid content, as C22-C24 SM, GalCer and ST were replaced by C18 variants. This led to fewer myelinated axons at 6 weeks of age, rapid myelin loss, and death between 4 and 6 months. Denatured myelin basic protein and a distinct population of phagocytic microglia were evident from 6 weeks of age, indicating that microglia sensed and responded to the myelin lipid defect soon after developmental myelination.

## Methods

### Generation of CerS2^fl/fl^ mice

Mice with loxP sites flanking exons 2 to 11 of the *CerS2* gene were generated by TaconicArtemis, Köln, Germany (Figure 2A). Mouse genomic fragments were sub-cloned using the RP23 BAC library. A neomycin resistance marker, flanked by FRT sites, was inserted into intron 1 and a puromycin selection marker, flanked by F3 sites, downstream of exon 11. The construct duplicated the 3’ untranslated region overlapping the *CerS2* and *Setdb1* genes, leaving the 3’ untranslated region of the *Setdb1* gene intact after Flp- and Cre-mediated deletion events. Linearized DNA vector was electroporated into C57BL/6N embryonic stem cells. Puromycin selection (1 μg/mL) and G418 selection (200 μg/mL) started on day 2, and counter-selection with Gancyclovir (2 μM) started on day 5 after electroporation. Correctly targeted clones were isolated on day 8 and injected into blastocysts isolated from super-ovulated Balb/c mice at 3.5 days post coitum. Flp-mediated removal of the neomycin and puromycin selection markers resulted in the floxed *CerS2* allele, confirmed by Southern blotting. PCR on genomic DNA using primers CAGCACCAAGACTCATCACC and CAAAACCCAGTCCGAGAAGC that span the 5’ loxP site yielded a 270 bp band for the wild-type and a 389 bp band for the floxed *CerS2* allele (Figure 2B). These mice were backcrossed to C57BL/6N for at least 8 generations prior to import into Australia, where they were maintained on a C57BL/6J background.

**Figure 2.**
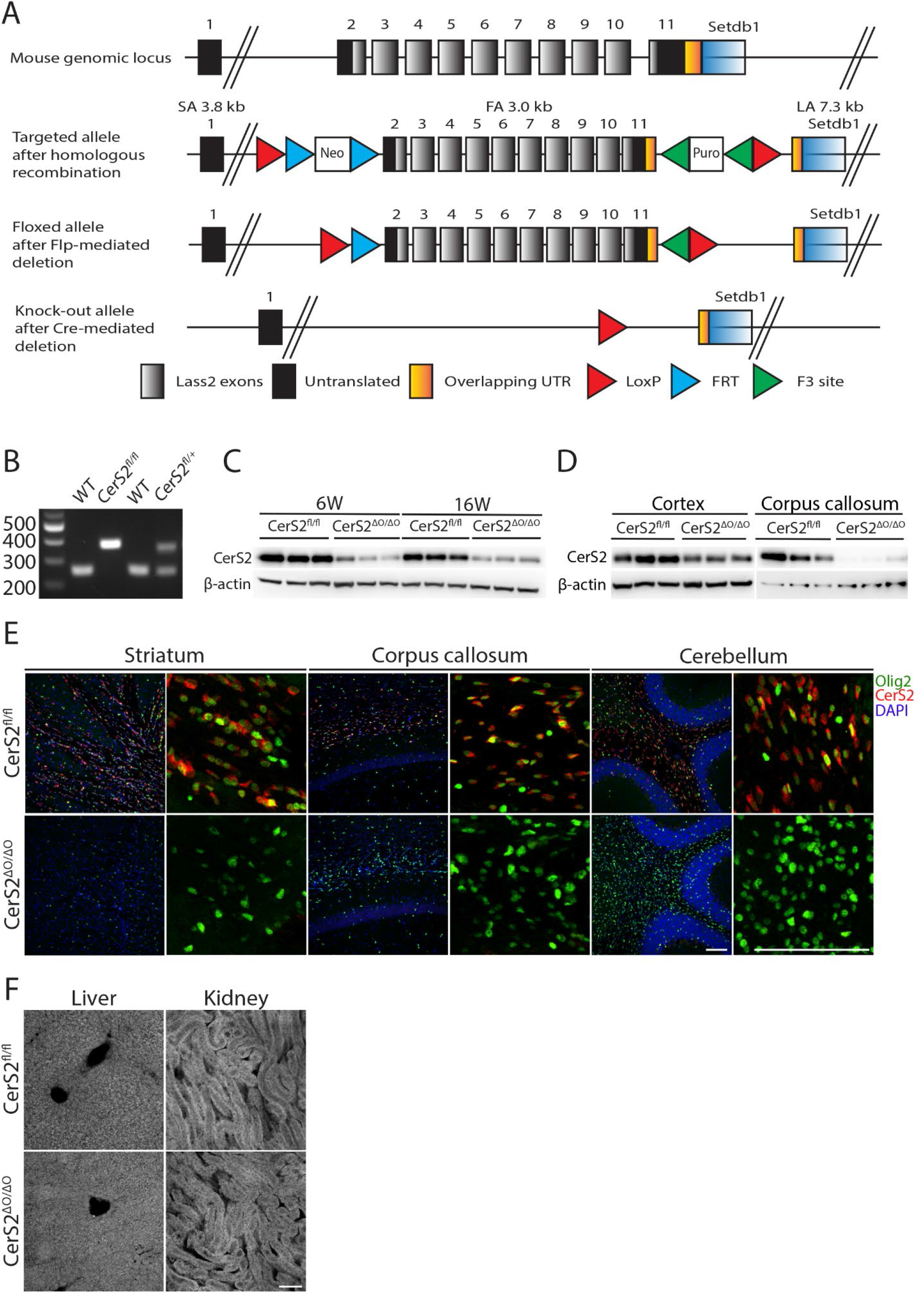
Generation of mice lacking CerS2 in oligodendrocytes. (A) Schematic for Cre-*lox*P-mediated deletion of exons 2 to 11 of the *CERS2* gene. (B) PCR genotyping assay for the floxed (380 bp) and wild-type *CERS2* allele (270 bp). (C-D) Western blots for CerS2 in (C) half-brain homogenates of 6- and 16-week-old (6W and 16W) male CerS2^fl/fl^ and CerS2^ΔO/ΔO^ mice, and (D) dissected cortex and corpus callosum of 6W female mice. (E) Immunofluorescent staining for CerS2 (red) and Olig2 (green) in the striatum, corpus callosum and cerebellum of 6W CerS2^fl/fl^ and CerS2^ΔO/ΔO^ mice. Lower magnification images include DAPI nuclear stain (blue). Scale bar: 100 µm. (F) Immunofluorescent staining for CerS2 in liver and kidneys.

### Generation and housing of conditional CerS2 knockout mice

To delete CerS2 in myelinating cells, mice homozygous for the floxed CerS2 allele (CerS2^fl/fl^) were bred to mice with heterozygous expression of Cre recombinase from the *CNP* gene locus (CNP-Cre) (Lappe-Siefke et al., 2003), which were also maintained on a C57BL/6J background. For experimental studies, CerS2^fl/fl^.CNP^cre/+^ mice were compared to CerS2^fl/fl^.CNP^+/+^ control littermates, resulting from CerS2^fl/fl^.CNP^cre/+^ × CerS2^fl/fl^.CNP^+/+^ breeding pairs. Male mice were used for all experiments, except where explicitly stated.

Experiments were conducted in accordance with the Australian code of practice for the care and use of animals for scientific purposes and approved by the University of Sydney animal ethics committee (#2017/1242). Mice were housed with food and water *ad libitum* on a 12-hour light/dark cycle. Following motor function tests, mice were anaesthetised with isofluorane, trans-cardially perfused with cold saline, and brains were removed. One hemisphere was frozen in liquid nitrogen for biochemical analyses. The other hemisphere was fixed in 4% paraformaldehyde (PFA, Sigma-Aldrich #2890) overnight at 4°C, followed by cryoprotection with 30% sucrose in phosphate buffered saline (PBS), then stored at −80°C.

### Motor function tests

A Rotarod apparatus (IITC Life Science, #755) was used to assess coordination and fatigue resistance (Carter et al., 2001). The rotarod started at 4 rpm and accelerated to 40 rpm over 60 s. The rotational speed and time at which each mouse fell off the rotarod was recorded. To measure forelimb grip strength, each mouse was allowed to grip a bar connected to a force transducer with both paws, and gently pulled away by the base of the tail while parallel to the ground. Similarly, the mouse was allowed to grip with all 4 paws on a grid to measure both fore- and hindlimb grip strength. The force (in Newtons) taken to ply the mouse away from the apparatus was recorded. The balance beam apparatus (Carter et al., 2001) was used to assess fine motor function and balance coordination. Each animal had to traverse an elevated, exposed, narrow 60 cm rod to reach the shelter of the goal box. Performance on the balance beam was quantified as the time taken to reach the goal, and the number of times their paw slipped off the beam. Mice were allowed to acclimatise to the goal box for 2 min and were trained on the apparatus by progressively increasing the starting distance away from the goal box. For all tests, each animal was tested 3 times. The average measurement for each mouse is reported.

### Immunofluorescence and histology staining and microscopy

Brains were cryosectioned at 40 μm along the sagittal plane and stored in cryoprotectant (25% ethylene glycol, 25% glycerol, 50% PBS) at −30 °C until use. Two to three sections per mouse were used for each stain. Sections were incubated in sodium citrate antigen retrieval buffer (10 mM, pH 6.0, 0.01% Tween 20) at 70°C for 10 min, blocked in PBS with 0.1% Tween-20 (PBST), 5% normal goat serum, and 0.1% bovine serum albumin (BSA) at RT for 2 h, and incubated overnight at 4°C with primary antibodies (Table 1) diluted in blocking solution. Sections were incubated in secondary antibodies (1:500) diluted in blocking solution for 2 h, counterstained with 1 μg/mL diamidino-2-phenylindole dihydrochloride (DAPI) and cover-slipped with ProLong Glass antifade (Life Technologies #P36980) before imaging. For Olig2, goat serum was excluded and BSA was increased to 5% for the required incubation steps. For every set of staining, one additional section was incubated in the absence of primary antibody as a negative control.

**Table 1.** Primary antibodies used for immunohistochemistry (IHC) and western blotting (WB).

| Primary antibody | Supplier | Dilution | Cat. # | RRID |
| --- | --- | --- | --- | --- |
| Rabbit anti-ASPA | Abcam | 1:1000 (IHC) | ab223269 | None found |
| Rabbit anti-CerS2 | Thermo Fisher | 1:1000<br>(WB/IHC) | PA5-55560 | AB_2643295 |
| Goat anti-Olig2 | R&D Systems | 1:1000 (IHC) | AF2418 | AB_2157554 |
| Rabbit anti-MBP | Abcam | 1:1000 (WB) | ab40390 | AB_1141521 |
| Rabbit anti-denatured MBP | Merck | 1:1000 (IHC) | ab5864 | AB_2140351 |
| Rabbit anti-MOG | Abcam | 1:1000 (WB) | ab32760 | AB_2145529 |
| Rabbit anti-PLP | Abcam | 1:1000 (WB) | ab28486 | AB_776593 |
| Mouse anti-βIII-tubulin | BioLegend | 1:1000 (WB) | 801202 | AB_10063408 |
| Rabbit anti-β-actin | Abcam | 1:5000 (WB) | ab8227 | AB_2305186 |
| Rabbit anti-APP (CT695) | Thermo Fisher | 1:1000 (IHC) | 512700 | AB_2533902 |
| Rabbit Anti-IBA1 | Abcam | 1:1000 (IHC) | ab178847 | AB_2832244 |
| Mouse anti-GFAP (GA5) | Cell Signalling Technology | 1:1000 (IHC) | 3670S | AB_561049 |

Luxol fast blue staining was performed as described (Song et al., 2021).

CerS2 and Olig2 staining was imaged on a Nikon C2 confocal microscope. All other fluorescent and histological staining was imaged with a Zeiss Axioscan slide scanner or an Olympus VS-120 slide scanner. Fiji image analysis software (ImageJ) was used to quantify ASPA-positive cells, APP puncta and the area of GFAP and IBA1 staining within defined white matter regions. Zeiss Zen Blue lite v3.4 software was used to acquire the mean intensity value of dMBP staining in the regions of interest.

### Western blotting

Frozen brain hemispheres were homogenised at 4°C in 20 mM Hepes pH 7.4, 5 mM NaF, 2 mM Na_3_VO_4_, 10 mM KCl and Complete mini EDTA-free protease inhibitor cocktail (Sigma #11836170001), using a Biospec mini bead beater with acid-washed glass beads (425-600 um). Cell debris was cleared by centrifugation at 1000×g for 15 min at 4°C and protein concentration was determined by bicinchoninic acid (BCA) assay (ThermoFisher Scientific #23225). Protein lysates were resolved on Bolt™ 4-12% Bis-Tris Plus gels (ThermoFisher Scientific #NW04125BOX) and transferred to polyvinylidene fluoride membranes. Membranes were blocked with 5% skim milk powder in Tris-buffered saline and 0.1% Tween 20 (TBST) for 1 h at RT, then incubated overnight with primary antibody diluted in 3% BSA. Membranes were washed in TBST thrice, incubated with horse radish peroxidase conjugated secondary antibody in blocking buffer for 1 h, washed in TBST three times, then imaged with ECL chemiluminescence reagent (EMD Millipore) on a Bio-Rad ChemiDoc Touch. Membranes were incubated in stripping buffer (15g/L glycine, 0.1% SDS, 1% Tween 20, pH 2.2) before re-probing for β-actin to control for protein loading. Bio-Rad Image Lab software (v6.0.1) was used for densitometry.

### Untargeted lipidomic analysis

Lipids were extracted from crude brain homogenates (∼1 mg homogenate protein) using a two-phase procedure with methyl-tert-butyl ether (MTBE)/methanol/water (10:3:2.5, v/v/v) (Couttas et al., 2020). The following internal standards were added to each sample: 2 nmole each of d18:1/12:0 SM, d18:1/12:0 ST, d18:1/12:0 glucosylceramide (GluCer), d18:1/17:0 ceramide, d17:1 sphingosine, d17:1 S1P, 17:0/17:0 phosphatidylethanolamine (PE), 17:0/17:0 phosphatidylserine (PS), 17:0/17:0 phosphatidylglycerol (PG), 17:0/17:0 phosphatidic acid (PA), 18:1/15:0-d7 diacylglycerol (DG), 17:0/17:0/17:0 triacylglycerol (TG), 14:0/14:0/14:0/14:0 cardiolipin (CL), 17:0 lysophosphatidic acid (LPA), 17:1 lysophosphatidylethanolamine (LPE), 17:1 lysophosphatidylserine (LPS), 18:1-d7 monoacylglycerol (MAG), 16:0-d3 carnitine, and 5 nmole 19:0/19:0 phosphatidylcholine (PC). TG internal standard was from Cayman Chemical. All other lipid standards were from Avanti Polar Lipids, distributed by Sigma Aldrich, Australia.

Untargeted lipidomic analysis was performed on a ThermoFisher Q-Exactive HF-X mass spectrometer with a heated electrospray ionization (HESI) probe and Vanquish HPLC system (Couttas et al., 2020). Extracts were resolved on a 2.1 × 100 mm Waters Acquity C18 HPLC column (1.7 µm pore size), using a 27 min binary gradient at 0.28 mL/min: 0 min, 80:20 A/B; 3 min, 80:20 A/B; 5.5 min, 55:45 A/B; 8 min, 35:65 A/B; 13 min, 15:85 A/B; 14 min, 0:100 A/B; 20 min, 0:100 A/B; 20.2 min, 70:30 A/B and 25 min, 70:30 A/B. Solvent A: 10 mM ammonium formate, 0.1% formic acid in acetonitrile:water (60:40); Solvent B: 10 mM ammonium formate, 0.1% formic acid in isopropanol:acetonitrile (90:10). Data was acquired in full scan/data-dependent MS^2^ (full scan resolution 60,000 FWHM, scan range 220–1600 *m/z*) in both positive and negative ionization modes. The ten most abundant ions in each cycle were subjected to MS^2^, with an isolation window of 1.1 *m/z*, collision energy 30 eV, resolution 15,000 FWHM, maximum integration time 50 ms and dynamic exclusion window 8 s. An exclusion list of background ions was used based on a solvent blank. An inclusion list of the [M+H]^+^ and [M-H]^−^ ions was used for all internal standards. LipidSearch software v4.2 (Thermo Fisher) was used for lipid annotation, chromatogram alignment, and peak integration from extracted ion chromatograms. Lipid annotation was based on precursor and product ions in both positive and negative ion mode. Individual lipids were expressed as ratios to an internal standard specific for each lipid class, then multiplied by the amount of internal standard added to produce a molar amount of each lipid per sample.

### Myelin isolation

Myelin isolation was performed as described (Larocca & Norton, 2007). Sagittal hemispheres were homogenised in 20 mM Hepes pH 7.4, 5 mM NaF, 2 mM Na_3_VO_4_, 10 mM KCl and EDTA-free protease inhibitor cocktail (Sigma #11836170001), and centrifuged at 10,000×g for 10 min. The pellet was resuspended in 20 mM Tris.HCl (pH 7.5), 2 mM Na_2_EDTA, 1 mM DTT, protease inhibitor cocktail, and 0.3 M sucrose, and layered over the same buffer containing 0.83 M sucrose, then centrifuged for 30 min at 75,000×g and 4°C. The white myelin interphase was collected into a new ultracentrifugation tube and exposed to an osmotic shock by vortexing in 20 mM Tris.HCl containing 2 mM Na_2_EDTA, 1 mM DTT and protease inhibitor cocktail. The myelin was pelleted by centrifugation for 15 min at 12,000×g and 4°C, and the osmotic shock was repeated. The myelin pellet was resuspended in 0.3 M sucrose, layered over 0.83 M sucrose and centrifuged at 75,000×g for 30 min, at 4°C. The osmotic shock procedure was repeated, and the myelin pellet was resuspended in 0.83 M sucrose, layered over 0.3 M sucrose, and centrifuged at 75,000×g for 30 min. The interphase was subjected to a final osmotic shock, pelleted at 75,000×g for 15 min, resuspended in Hepes buffer, and protein concentration was determined using the BCA assay.

### Targeted lipidomic analysis and GalCer/GluCer isomer separation

Lipids were extracted from ∼50 μg isolated myelin protein or ∼500 μg liver homogenate and analysed by multiple reaction monitoring on a TSQ Altis triple quadrupole mass spectrometer, as described (Song et al., 2021). An internal standard for each lipid class was included, as described above. Lipids were quantified relative to external standard curves created with a synthetic standard for each lipid class, expressed as a ratio to the relevant internal standard for that lipid class. Lipid standards were purchased from Avanti Polar Lipids.

Separation of GalCer from GluCer was achieved using a 2.1 x 150 mm Agilent InfinityLab Poroshell 120 HILIC-Z HPLC column (2.7 µm pore size) on a Shimazu Nexera HPLC coupled to a Sciex 6500+ QTRAP mass spectrometer. The HPLC solvent was 97% acetonitrile, 2% methanol, 0.75% water, 0.25% formic acid, and 2.5 mM ammonium formate. Lipid extracts were reconstituted in this solvent prior to HPLC. Precursor *m/z* was the relevant [M+H]^+^ ion, and product *m/z* was 264.3. Run time was 20 min, and flow rate 0.1 mL/min. SciexOS software was used for lipid annotation, chromatogram alignment, and peak integration.

### Transmission Electron Microscopy (TEM)

Animals were transcardially perfused with sterile saline, followed by 2.5% glutaraldehyde, 1% PFA in 0.1M PBS. Brains were post-fixed in the same fixative at 4°C overnight. The first two millimetres of the sagittal midline of the right hemisphere were selected for EM processing. For analysis of the corpus callosum, the first millimetre from Bregma −1.1 to −3.2mm containing the corpus callosum was micro-dissected, while for analysis of the striatum radiatum, the second millimetre from the sagittal midline containing the hippocampus was used. Selected tissue regions were placed in Kanovsky’s buffer overnight, then washed in 0.1M sodium cacodylate and embedded in the sagittal plane in epoxy resin. Semi-thin (0.5-1µm) sections were collected on glass slides and stained with 1% toluidine blue to identify regions of interest. Subsequent ultrathin (0.1µm) sections were collected on 3×3mm copper grids and contrasted with heavy metals. Contrasted and un-contrasted ultrathin sections were viewed using a JEOL JEM-1400Flash TEM. Six distinct fields of view per mouse were imaged at 15,000x magnification and captured as 3×3 tile scans using the JEOL integrated software and a high-sensitivity sCMOS camera (JEOL Matataki Flash). G-ratios were calculated as the diameter of the axon lumen divided by the diameter of the lumen plus myelin sheath (Song et al., 2021). These measurements were taken from 3 mice per genotype, and 220 - 500 axons per mouse, using ImageJ software.

### Whole-brain flow cytometry

Brains were processed into single cell suspensions for spectral cytometry, as previously described (Spiteri et al., 2022; Spiteri et al., 2021). Briefly, brain cells were isolated using a 30%/80% Percoll gradient after homogenizing tissue on a gentleMACS dissociator (Miltenyi Biotec). Single cell suspensions were then incubated with purified anti-CD16/32 (Biolegend, RRID: AB_312801) and Zombie UV Fixable Viability kit (Biolegend, USA) before staining with a cocktail of fluorescently-labelled antibodies. Fluorochrome-conjugated antibodies used for surface staining included: anti-CD11b (M1/70, Biolegend), anti-CD11c (HL3, BD Biosciences), anti-B220 (RA3-6B2, Biolegend and BD Biosciences), anti-CD8α (53-6.7, BD Biosciences), anti-CX3CR1 (SA011F11, Biolegend), anti-I-A/I-E (M5/114.15.2, Biolegend), anti-Ly6C (HK1.4, Biolegend), anti-Ly6G (1A8, Biolegend), anti-F4/80 (BM8, Biolegend), anti-CD4 (RM4-5, Biolegend), anti-P2RY12 (S16007D, Biolegend), anti-CD64 (X54-5/7.1, Biolegend), anti-CD45 (30-F11, Biolegend), anti-CD3ε (145-2C11, Biolegend), anti-NK1.1 (PK136, Biolegend), TMEM119 (106-6, Abcam) and anti-CD86 (GL-1, Biolegend). Cells were washed twice and fixed in fixation buffer (Biolegend). For intracellular staining, cells were then incubated with Cytofix/Cytoperm (BD Biosciences) before staining with anti-CD68 (FA-11, Biolegend) and anti-CD206 (C068C2, Biolegend). Cells were washed twice and subsequently analysed on a 5-laser Aurora Spectral Cytometer (Cytek Biosciences). Acquired data was analysed in FlowJo (v10.8, BD Biosciences). Channel values (CSV) were exported from total microglial populations in FlowJo. Clustering and dimensionality reduction were performed using FlowSOM (Van Gassen et al., 2015) and Uniform Manifold Approximation and Projection (UMAP) (Becht et al., 2018) on total microglia populations in RStudio (1.1.453 or 1.4.1717) using *Spectre* (Ashhurst et al., 2021) with default settings. Microglial FlowSOM clusters were identified using manual gating in FlowJo (Supplementary Fig. 3). Median fluorescent intensity (MFI) signals were exported from these microglial subsets in FlowJo to generate a heatmap in RStudio (1.2.1335) using the R package, *pheatmaps*.

### Statistical analysis

All statistical analyses were performed with GraphPad Prism. Results testing the effect of both genotype and age were analysed by two-way ANOVA, followed by Bonferroni’s post-hoc test. Results testing the effect of genotype at a single age were analysed by t-tests. Data that were not normally distributed were analysed by the Mann-Whitney test. P values < 0.05 were considered significant. Lipidomic data was analysed by unpaired t-tests, applying the Benjamini, Krieger and Yekutieli method to adjust for false discovery rate (*q* < 0.01).

### Data Availability

The untargeted lipidomic data associated with this manuscript is available as Supplementary Table 1. All other data that support the findings of this study are available from the corresponding author upon reasonable request.

## Results

### Deletion of *CERS2* in oligodendrocytes

Targeted deletion of CerS2 was facilitated by introducing loxP sites flanking exons 2 to 11 of the *CERS2* gene (Fig. 2A-B). Mice homozygous for the floxed CerS2 allele (CerS2^fl/fl^) were bred to the CNP-Cre line, resulting in CerS2^fl/fl^ mice heterozygous for the CNP-Cre allele, which are hereafter referred to as CerS2^ΔO/ΔO^. CerS2^fl/fl^ littermates without CNP-Cre were used as controls. CerS2 protein was notably reduced in whole brain homogenates of 6- and 16-week-old male CerS2^ΔO/ΔO^ mice (Fig. 2C). Western blotting on dissected white matter (corpus callosum) from 6-week-old females showed almost complete loss of CerS2 protein, whereas CerS2 levels were only partially reduced in cortex (Fig. 2D), supporting the oligodendrocyte-specific nature of the deletion. CerS2 immunostaining co-localised with the oligodendrocyte lineage marker Olig2 in white matter (Fig. 2E). This oligodendrocyte CerS2 staining was lost in CerS2^ΔO/ΔO^ mice (Fig. 2E), whereas CerS2 immunoreactivity in liver and kidney was unaffected (Fig 2F).

### Myelin lipid acyl chain length is decreased in CerS2^ΔO/ΔO^ mice

To understand how CerS2 deletion in oligodendrocytes affects the brain lipidome, we performed untargeted lipidomic analysis on brain homogenates of 6-week-old male mice. A total of 532 phospholipids, sphingolipids, and neutral lipids were identified, based on both precursor and product ions, and quantified relative to internal standards (Supplementary Table 1). Of these, 20% (107) were differentially expressed between CerS2^fl/fl^ and CerS2^ΔO/ΔO^ mice at a false discovery rate-adjusted *p* value of 0.01 (Fig. 3A and Table 2). The most heavily affected lipid classes were the major myelin sphingolipids HexCer (42/55 species differentially expressed), and ST (10/10 differentially expressed), as well as ceramides (11/27 differentially expressed). Together with SM (7/28 differentially expressed), these sphingolipids made up two thirds of the most significantly regulated lipids, with the remaining third comprised of glycerophospholipids. The most significantly decreased lipids in CerS2^ΔO/ΔO^ mice were HexCer and ST with N-acyl chain lengths greater than 20 carbons, whereas the most significantly increased were HexCer with C16 or C18 N-acyl chains (Table 1), indicating that the most significant change to the brain lipidome was a shift from very long chain (C22-C24) to long chain (C16-C18) sphingolipids, as predicted for CerS2 deletion.

**Figure 3.**
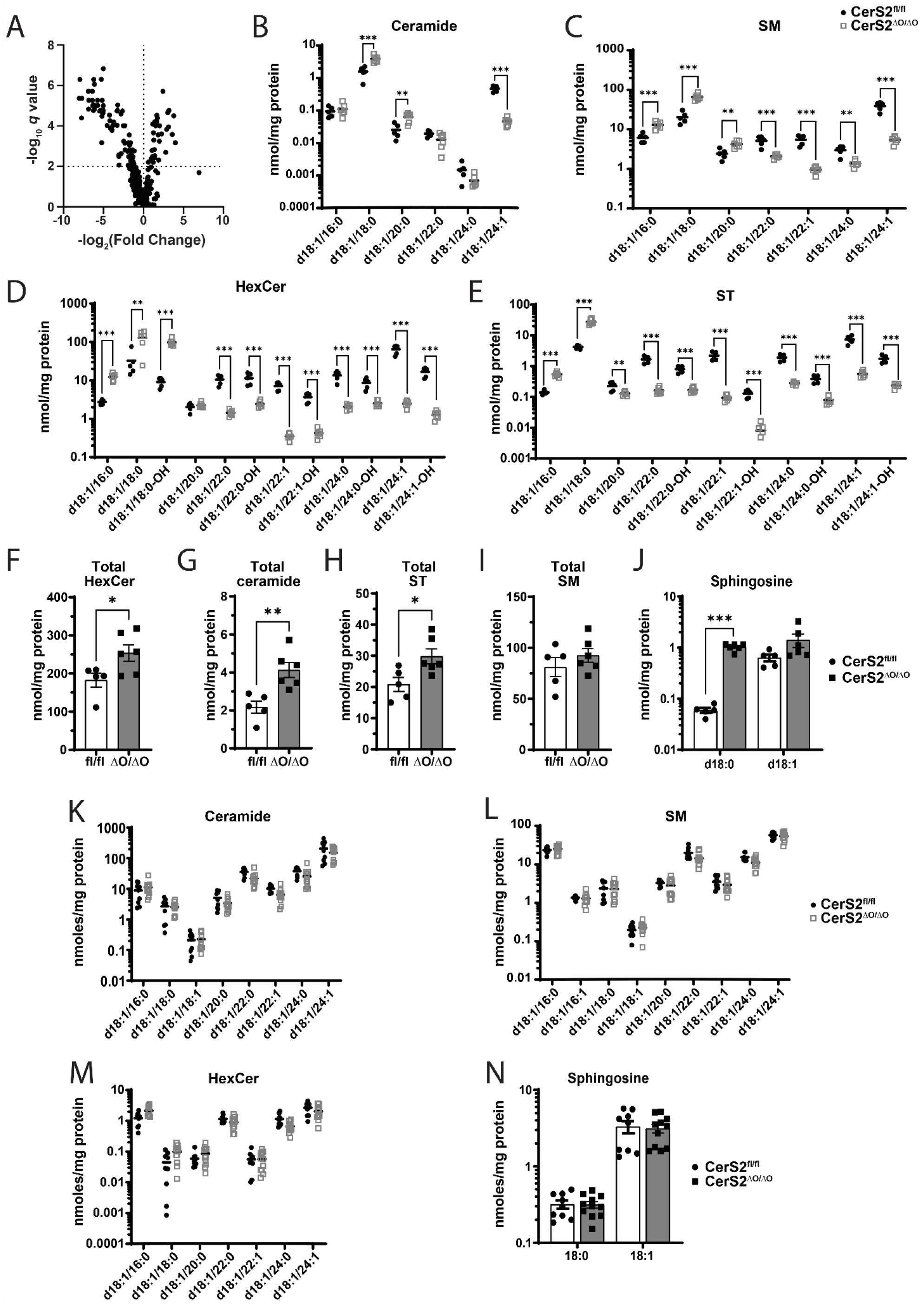
Increased C16-C18 and decreased C22-C24 sphingolipids in myelin from 6W CerS2^ΔO/ΔO^ mice. (A) Volcano plot showing mean fold change and corresponding *q* value (*p* value adjusted for false discovery rate) for changes in lipid species in 6-week-old male CerS2^ΔO/ΔO^ mice (n = 7) relative to their CerS2^fl/fl^ littermates (n = 6). (B-J) Sphingolipid levels in isolated myelin from female CerS2^fl/fl^ (circles, n = 5) and CerS2^ΔO/ΔO^ (open squares, n = 6) mice. Individual (B) ceramide, (C) sphingomyelin (SM), (D) Hexosylceramide (HexCer), and (E) sulfatide (ST) species; total (F) HexCer, (G) ceramide, (H) ST and (I) SM levels; and (J) dihydrosphingosine (d18:0) and sphingosine (d18:1) levels. (K) Ceramide, (L) SM, (M) HexCer, and (N) sphingosine levels in liver samples from 6-week-old CerS2^fl/fl^ (circles, n = 4 males, n = 5 females) and CerS2^ΔO/ΔO^ (open squares, n = 5 males, n = 6 females) mice. T-tests adjusted for multiple comparisons (false discovery rate 1%) were used to compare levels of individual lipids between the two genotypes (B-E and J-N); ** *p*<0.01, *** *p*<0.001. Unadjusted t-tests were used to compare lipid totals (F-I).

**Table 2.** Lipids differing significantly in abundance between CerS2^ΔO/ΔO^ and CerS2^fl/fl^ brains. Lipids were quantified in brain homogenates of 6 week old male mice by untargeted lipidomic profiling. Lipids differing significantly in abundance between the genotypes were identified by t-tests adjusted for multiple comparisons (*q* < 0.01).

| Lipid and ion adduct | Mean CerS2<sup>fl/fl</sup> | SD CerS2<sup>fl/fl</sup> | Mean CerS2<sup>ΔO/ΔO</sup> | SD CerS2<sup>ΔO/ΔO</sup> | T ratio | *p* value | *q* value |
| --- | --- | --- | --- | --- | --- | --- | --- |
| ST(d18:2_24:1)+H | 4385.54 | 1540.04 | 133.92 | 36.63 | 20.83 | 3.00E-10 | 1.49E-07 |
| ST(d18:1_24:1)+H | 49891.33 | 18253.91 | 1287.83 | 441.15 | 17.39 | 2.40E-09 | 5.05E-07 |
| PC(42:7e)+H | 332.99 | 97.97 | 1.34 | 0.81 | 16.77 | 3.50E-09 | 5.05E-07 |
| Hex1Cer(d18:1_24:3)+H | 948.12 | 672.72 | 12.54 | 5.23 | 16.32 | 4.70E-09 | 5.05E-07 |
| Hex1Cer(d18:2_24:0+O)+H | 4645.54 | 3333.27 | 65.85 | 32.88 | 15.17 | 1.01E-08 | 8.75E-07 |
| LPC(24:0)+HCOO | 26.57 | 5.35 | 141.08 | 31.12 | 13.81 | 2.71E-08 | 1.93E-06 |
| Hex1Cer(d18:0_24:1)+H | 727.77 | 773.77 | 6.02 | 2.65 | 13.62 | 3.14E-08 | 1.93E-06 |
| Hex1Cer(d18:1_24:2)+H | 12588.75 | 9489.57 | 149.22 | 104.66 | 13.45 | 3.58E-08 | 1.93E-06 |
| Hex1Cer(d18:1_25:1+O)+H | 72.06 | 76.95 | 0.34 | 0.26 | 13.49 | 9.66E-08 | 4.25E-06 |
| Hex1Cer(d18:1_25:2)+H | 159.82 | 138.82 | 0.61 | 0.51 | 13.46 | 9.83E-08 | 4.25E-06 |
| Hex1Cer(d18:1_23:0+O)+H | 732.96 | 501.28 | 20.11 | 9.41 | 11.82 | 1.36E-07 | 5.36E-06 |
| ST(d18:0_24:1)+H | 3451.12 | 2028.35 | 29.97 | 21.68 | 12.78 | 1.61E-07 | 5.42E-06 |
| Hex1Cer(d18:1_25:1)+H | 310.05 | 292.46 | 3.67 | 2.45 | 11.59 | 1.67E-07 | 5.42E-06 |
| Hex1Cer(d18:1_24:1)+H | 22734.37 | 19500.37 | 422.18 | 304.11 | 11.53 | 1.76E-07 | 5.42E-06 |
| PE(20:1_18:1)+H | 1885.19 | 809.35 | 199.71 | 76.87 | 10.82 | 3.36E-07 | 9.11E-06 |
| Hex1Cer(d18:2_22:1)+H | 519.85 | 366.44 | 6.99 | 5.08 | 10.81 | 3.37E-07 | 9.11E-06 |
| Hex1Cer(d18:1_23:1)+H | 2134.31 | 1847.74 | 39.68 | 28.79 | 10.67 | 3.86E-07 | 9.82E-06 |
| Hex1Cer(d18:1_23:1+O)+H | 165.37 | 140.34 | 1.99 | 1.74 | 11.52 | 4.28E-07 | 9.87E-06 |
| ST(d42:2+O)+H | 4532.43 | 2352.83 | 138.81 | 145.38 | 10.55 | 4.34E-07 | 9.87E-06 |
| Hex1Cer(d18:1_24:1+O)+H | 4681.57 | 3426.56 | 39.74 | 44.39 | 10.08 | 6.82E-07 | 1.47E-05 |
| Hex1Cer(d18:1_22:2)+H | 1224.55 | 815.37 | 17.29 | 14.78 | 10.03 | 7.18E-07 | 1.48E-05 |
| Hex1Cer(d44:2+O)+H | 84.33 | 82.53 | 2.18 | 0.99 | 9.86 | 8.55E-07 | 1.68E-05 |
| PC(17:1_18:1)+H | 1100.84 | 252.50 | 113.85 | 75.42 | 9.80 | 9.00E-07 | 1.69E-05 |
| Hex1Cer(d18:1_26:2)+H | 216.24 | 193.93 | 2.82 | 2.53 | 9.70 | 1.00E-06 | 1.71E-05 |
| Hex1Cer(d18:1_18:0+O)+H | 1049.97 | 642.03 | 9419.91 | 3735.38 | 9.69 | 1.02E-06 | 1.71E-05 |
| Hex1Cer(d18:1_22:0)+H | 1098.60 | 991.82 | 33.52 | 21.93 | 9.67 | 1.03E-06 | 1.71E-05 |
| Hex1Cer(d18:1_23:0)+H | 275.05 | 289.95 | 6.04 | 5.74 | 9.58 | 1.14E-06 | 1.79E-05 |
| PC(40:4e)+H | 429.92 | 165.99 | 43.73 | 20.42 | 9.56 | 1.16E-06 | 1.79E-05 |
| SM(d44:5)+H | 1144.69 | 260.76 | 183.44 | 83.78 | 9.49 | 1.25E-06 | 1.86E-05 |
| Hex1Cer(d18:1_25:0+O)+H | 54.64 | 46.29 | 1.45 | 1.04 | 9.42 | 1.34E-06 | 1.93E-05 |
| Hex1Cer(d18:1_18:1)+H | 3390.77 | 2157.79 | 27364.71 | 10971.31 | 9.26 | 1.59E-06 | 2.21E-05 |
| Hex1Cer(d18:1_24:0)+H | 1621.41 | 1624.88 | 47.82 | 31.34 | 9.19 | 1.71E-06 | 2.31E-05 |
| Cer(d41:2)+HCOO | 94.54 | 14.41 | 13.42 | 6.19 | 9.07 | 1.94E-06 | 2.55E-05 |
| Hex1Cer(d18:1_22:1)+H | 10101.18 | 7831.03 | 531.61 | 362.88 | 8.92 | 2.29E-06 | 2.92E-05 |
| Cer(d18:0_18:0)+HCOO | 180.64 | 34.70 | 564.30 | 130.37 | 8.87 | 2.41E-06 | 2.98E-05 |
| Hex1Cer(d18:0_18:0)+H | 204.96 | 186.74 | 2764.73 | 1238.04 | 8.79 | 2.64E-06 | 3.17E-05 |
| ST(d18:1_24:2)+H | 2948.60 | 1517.68 | 80.67 | 13.44 | 13.54 | 2.81E-06 | 3.29E-05 |
| PG(18:0_20:4)-H | 231.91 | 54.80 | 763.26 | 187.87 | 8.52 | 3.59E-06 | 4.06E-05 |
| Hex1Cer(d18:1_23:2)+H | 272.17 | 236.22 | 1.26 | 1.93 | 9.97 | 3.67E-06 | 4.06E-05 |
| Hex1Cer(d18:1_22:0+O)+H | 3207.36 | 2604.14 | 174.76 | 129.48 | 8.22 | 5.04E-06 | 5.44E-05 |
| Hex1Cer(d38:4)+H | 5704.68 | 1195.41 | 17670.39 | 5161.30 | 8.11 | 5.70E-06 | 6.01E-05 |
| PG(36:4)-H | 11.02 | 2.23 | 29.37 | 6.31 | 8.08 | 5.93E-06 | 6.10E-05 |
| ST(d18:1_24:0)+H | 15067.14 | 10368.01 | 1121.18 | 812.62 | 8.02 | 6.35E-06 | 6.39E-05 |
| PG(18:1_20:4)-H | 11.98 | 2.39 | 26.41 | 3.95 | 7.85 | 7.83E-06 | 7.70E-05 |
| SM(d42:2)+H | 7852.93 | 5687.34 | 793.61 | 386.10 | 7.68 | 9.60E-06 | 9.22E-05 |
| Hex1Cer(d18:1_24:0+O)+H | 3048.65 | 2729.29 | 130.18 | 93.26 | 7.65 | 9.92E-06 | 9.32E-05 |
| PE(18:0_20:1)+H | 1268.87 | 563.88 | 250.36 | 81.05 | 7.61 | 1.05E-05 | 9.65E-05 |
| Cer(d43:2)+HCOO | 16.66 | 5.92 | 1.16 | 0.81 | 8.06 | 1.11E-05 | 9.82E-05 |
| SM(d33:1)+H | 293.05 | 71.98 | 50.95 | 33.37 | 7.55 | 1.12E-05 | 9.82E-05 |
| PC(41:2)+H | 6.36 | 2.30 | 25.20 | 7.05 | 7.54 | 1.14E-05 | 9.82E-05 |
| Hex1Cer(d18:2_18:0+O)+H | 5.76 | 1.34 | 16.13 | 4.09 | 7.53 | 1.16E-05 | 9.85E-05 |
| Hex1Cer(d18:1_19:1)+H | 39.41 | 34.46 | 342.71 | 177.17 | 7.41 | 1.34E-05 | 1.11E-04 |
| Hex1Cer(d18:0_24:0)+H | 130.29 | 123.87 | 7.37 | 4.68 | 7.29 | 1.56E-05 | 1.27E-04 |
| SM(d41:4)+H | 13.70 | 8.27 | 101.25 | 52.55 | 7.16 | 1.85E-05 | 1.48E-04 |
| Hex1Cer(d18:0_22:0)+H | 210.92 | 223.74 | 13.73 | 7.47 | 7.06 | 2.10E-05 | 1.65E-04 |
| SM(d35:1)+H | 269.63 | 80.98 | 624.92 | 109.47 | 6.84 | 2.80E-05 | 2.16E-04 |
| Hex1Cer(d18:1_18:1+O)+H | 49.64 | 32.11 | 232.75 | 45.45 | 6.73 | 3.24E-05 | 2.46E-04 |
| Hex1Cer(t18:0_18:0)+H | 2.89 | 2.77 | 34.59 | 10.62 | 6.67 | 3.51E-05 | 2.62E-04 |
| PC(15:0_18:1)+H | 5304.51 | 1000.36 | 1538.39 | 874.50 | 6.63 | 3.69E-05 | 2.70E-04 |
| Hex1Cer(d18:1_16:0)+H | 204.73 | 108.14 | 722.49 | 231.08 | 6.61 | 3.80E-05 | 2.74E-04 |
| LPC(24:1)+HCOO | 6.55 | 1.53 | 15.57 | 4.06 | 6.20 | 6.72E-05 | 4.76E-04 |
| Hex1Cer(d18:1_18:0)+H | 3213.54 | 2191.03 | 12544.07 | 4467.91 | 6.07 | 8.04E-05 | 5.61E-04 |
| ST(d18:1_24:0+O)+H | 3632.19 | 2088.11 | 645.41 | 485.04 | 6.05 | 8.26E-05 | 5.66E-04 |
| PC(16:0_23:1)+H | 499.67 | 70.97 | 159.32 | 92.32 | 5.93 | 9.91E-05 | 6.64E-04 |
| Hex1Cer(d18:1_17:0)+H | 5.23 | 7.84 | 81.79 | 29.41 | 6.58 | 1.01E-04 | 6.64E-04 |
| Hex1Cer(d18:0_23:0)+H | 28.05 | 31.71 | 0.75 | 0.70 | 5.91 | 1.01E-04 | 6.64E-04 |
| PE(18:1p_22:1)+H | 605.40 | 271.51 | 165.77 | 61.70 | 5.89 | 1.04E-04 | 6.70E-04 |
| ST(d18:1_22:0+O)+H | 4155.67 | 2435.19 | 824.81 | 511.25 | 5.85 | 1.12E-04 | 7.09E-04 |
| PE(18:0_18:1)+H | 8777.99 | 3714.43 | 2626.28 | 893.57 | 5.82 | 1.16E-04 | 7.29E-04 |
| Cer(d18:1_24:1)+HCOO | 1710.26 | 504.97 | 510.79 | 224.55 | 5.68 | 1.42E-04 | 8.79E-04 |
| SM(d36:0)+H | 1074.75 | 522.12 | 2720.63 | 642.26 | 5.35 | 2.35E-04 | 1.43E-03 |
| PS(42:5e)-H | 47.90 | 13.61 | 18.49 | 7.67 | 5.29 | 2.56E-04 | 1.53E-03 |
| PE(16:0p_20:0)+H | 86.54 | 45.84 | 17.46 | 5.02 | 5.28 | 2.59E-04 | 1.53E-03 |
| Cer(d18:0_23:0)+HCOO | 15.69 | 4.03 | 4.60 | 2.64 | 5.26 | 2.67E-04 | 1.56E-03 |
| PG(18:1_18:1)-H | 64.84 | 17.36 | 133.65 | 24.86 | 5.24 | 2.79E-04 | 1.61E-03 |
| PS(40:4e)-H | 240.80 | 58.34 | 108.18 | 32.82 | 5.18 | 3.02E-04 | 1.72E-03 |
| SM(d18:1_24:2)+H | 491.72 | 236.16 | 149.28 | 58.62 | 5.17 | 3.11E-04 | 1.74E-03 |
| PE(39:2e)+H | 57.78 | 26.31 | 14.42 | 4.95 | 5.15 | 3.19E-04 | 1.77E-03 |
| Hex1Cer(d16:1_18:1)+H | 15.30 | 22.19 | 103.92 | 46.97 | 5.27 | 3.62E-04 | 1.98E-03 |
| ST(d18:0_24:0)+H | 982.12 | 756.58 | 141.51 | 122.46 | 4.96 | 4.26E-04 | 2.30E-03 |
| PE(18:1p_20:1)+H | 4363.04 | 2172.80 | 1505.25 | 568.50 | 4.89 | 4.83E-04 | 2.58E-03 |
| PS(18:1_22:1)-H | 660.58 | 115.48 | 231.28 | 187.93 | 4.86 | 5.03E-04 | 2.65E-03 |
| Hex1Cer(d18:1_26:1)+H | 171.48 | 183.95 | 19.96 | 15.24 | 4.84 | 5.19E-04 | 2.70E-03 |
| PC(40:0)+H | 132.47 | 29.99 | 346.30 | 166.95 | 4.78 | 5.70E-04 | 2.94E-03 |
| PG(38:3)-H | 10.40 | 3.30 | 22.69 | 5.25 | 4.75 | 6.04E-04 | 3.07E-03 |
| PC(15:0_20:3)+H | 717.21 | 173.41 | 275.22 | 156.78 | 4.73 | 6.24E-04 | 3.14E-03 |
| Hex1Cer(d18:1_21:1)+H | 138.87 | 133.88 | 16.40 | 17.66 | 4.71 | 6.44E-04 | 3.20E-03 |
| Cer(d41:1)+HCOO | 73.62 | 29.10 | 20.22 | 8.92 | 4.69 | 6.61E-04 | 3.24E-03 |
| Cer(d40:2)+HCOO | 331.65 | 78.01 | 132.06 | 59.81 | 4.67 | 6.85E-04 | 3.33E-03 |
| PE(18:1p_14:0)+H | 273.27 | 132.89 | 87.88 | 35.32 | 4.63 | 7.24E-04 | 3.48E-03 |
| Cer(d18:2_24:1)+HCOO | 197.69 | 27.16 | 501.62 | 283.17 | 4.58 | 7.93E-04 | 3.73E-03 |
| PC(44:4)+H | 362.92 | 75.31 | 131.64 | 69.92 | 4.58 | 7.94E-04 | 3.73E-03 |
| Cer(d18:0_22:0)+HCOO | 38.80 | 18.18 | 11.92 | 4.94 | 4.56 | 8.13E-04 | 3.78E-03 |
| LPC(22:0)+HCOO | 8.96 | 2.34 | 16.97 | 4.00 | 4.51 | 8.84E-04 | 4.07E-03 |
| PE(18:0p_20:1)+H | 2938.37 | 1390.57 | 1031.54 | 308.59 | 4.45 | 9.76E-04 | 4.44E-03 |
| Hex1Cer(d18:1_16:1)+H | 3.84 | 2.95 | 14.00 | 3.03 | 4.40 | 1.06E-03 | 4.79E-03 |
| Cer(d43:1)+HCOO | 11.78 | 5.09 | 3.81 | 2.21 | 4.33 | 1.18E-03 | 5.28E-03 |
| PE(18:2e_22:4)+H | 759.14 | 440.50 | 263.99 | 98.60 | 4.29 | 1.28E-03 | 5.61E-03 |
| PC(40:2e)+H | 100.35 | 16.76 | 40.42 | 29.28 | 4.29 | 1.29E-03 | 5.61E-03 |
| PC(42:1)+H | 1869.42 | 281.24 | 4315.17 | 2270.15 | 4.22 | 1.43E-03 | 6.18E-03 |
| PG(32:1)-H | 26.30 | 2.81 | 39.06 | 7.35 | 4.21 | 1.47E-03 | 6.29E-03 |
| PE(18:1_18:1)+H | 12762.44 | 6461.69 | 4746.90 | 2370.95 | 4.14 | 1.66E-03 | 7.01E-03 |
| Cer(d18:0_26:0)+HCOO | 21.23 | 3.68 | 9.13 | 5.88 | 4.11 | 1.72E-03 | 7.22E-03 |
| ST(d18:1_22:1)+H | 2323.24 | 1581.92 | 277.93 | 244.98 | 4.02 | 2.03E-03 | 8.43E-03 |
| PE(20:1_22:6)+H | 151.09 | 106.09 | 39.85 | 31.65 | 3.99 | 2.12E-03 | 8.73E-03 |
| PE(16:0_22:4)+H | 686.20 | 431.71 | 216.91 | 144.43 | 3.97 | 2.20E-03 | 8.99E-03 |
| PE(18:0_20:3)+H | 1236.85 | 590.82 | 506.49 | 198.01 | 3.93 | 2.34E-03 | 9.45E-03 |

To gain a more detailed understanding of how CerS2 deletion in oligodendrocytes affects the sphingolipid composition of myelin, targeted lipidomic analysis was performed on purified myelin from 6-week-old female mice. CerS2 deletion in oligodendrocytes led to loss of C24:1 ceramide, and a compensatory increase in C18:0 and C20:0 (Fig. 3B). These changes were more marked in the ceramide derivatives SM, HexCer, and ST, in which loss of C22-C24 and gain of C16-C18 species was clearly apparent (Fig. 3C-E). Increased C16-C20 variants produced an increase in total HexCer, ST and ceramide in myelin from CerS2^ΔO/ΔO^ relative to CerS2^fl/fl^ mice, despite reduced C22-C24 variants (Fig. 3F-H), whereas total SM was not significantly different (Fig. 3I). The ceramide precursor dihydrosphingosine was an order of magnitude higher in the myelin fraction of CerS2^ΔO/ΔO^ mice whereas sphingosine was not significantly different (Fig. 3J). Myelin cholesterol content did not differ between CerS2^fl/fl^ and CerS2^ΔO/ΔO^ mice (1577 ± 202 vs 1616 ± 140 nmoles/mg protein), and sphingolipid levels in the liver did not differ between the two genotypes (Fig 3K-N).

HexCer comprises both GalCer and GluCer, which are indistinguishable using LC-MS/MS with reverse-phase chromatography. Chemical analyses have previously established that > 99% HexCer in the brain is GalCer (Vanier & Svennerholm, 1975). Using hydrophilic interaction (HILIC) chromatography we confirmed that the increased C18 HexCer content of both brain homogenates and purified myelin was GalCer, not GluCer (Supplementary Fig. 1). C18 and C24:1 GluCer were undetectable in myelin.

### Myelin degeneration and early lethality in CerS2^ΔO/ΔO^ mice

In our initial characterisation of CerS2^ΔO/ΔO^ mice, sudden, unexpected deaths occurred, starting from 14-16 weeks of age and affecting 80% of the mice by 22 weeks (Fig. 4A). Spontaneous tonic seizures were also observed in CerS2^ΔO/ΔO^ mice. Since death is not an ethical endpoint, CerS2^ΔO/ΔO^ mice were subsequently kept up to 16 weeks of age. There was no difference in litter size (Fig 4B) or body weight (Fig. 4C) between CerS2^ΔO/ΔO^ mice and their CerS2^fl/fl^ littermates. However, pronounced myelin loss was apparent in luxol fast blue stained sections from 16-week-old CerS2^ΔO/ΔO^ mice (Fig. 4D), and levels of the myelin protein markers myelin basic protein (MBP), proteolipid protein (PLP), and myelin oligodendrocyte glycoprotein (MOG) were greatly reduced (Fig. 4E-F). Myelin staining (Fig. 4D) and protein levels (Fig. 4E-F) were not reduced in CerS2^ΔO/ΔO^ relative to CerS2^fl/fl^ mice at 6 weeks of age, apart from a small reduction in total PLP content. The neuronal marker βIII-tubulin was very modestly reduced in 16-week-old CerS2^ΔO/ΔO^ mice (Fig. 4E-F).

**Figure 4:**
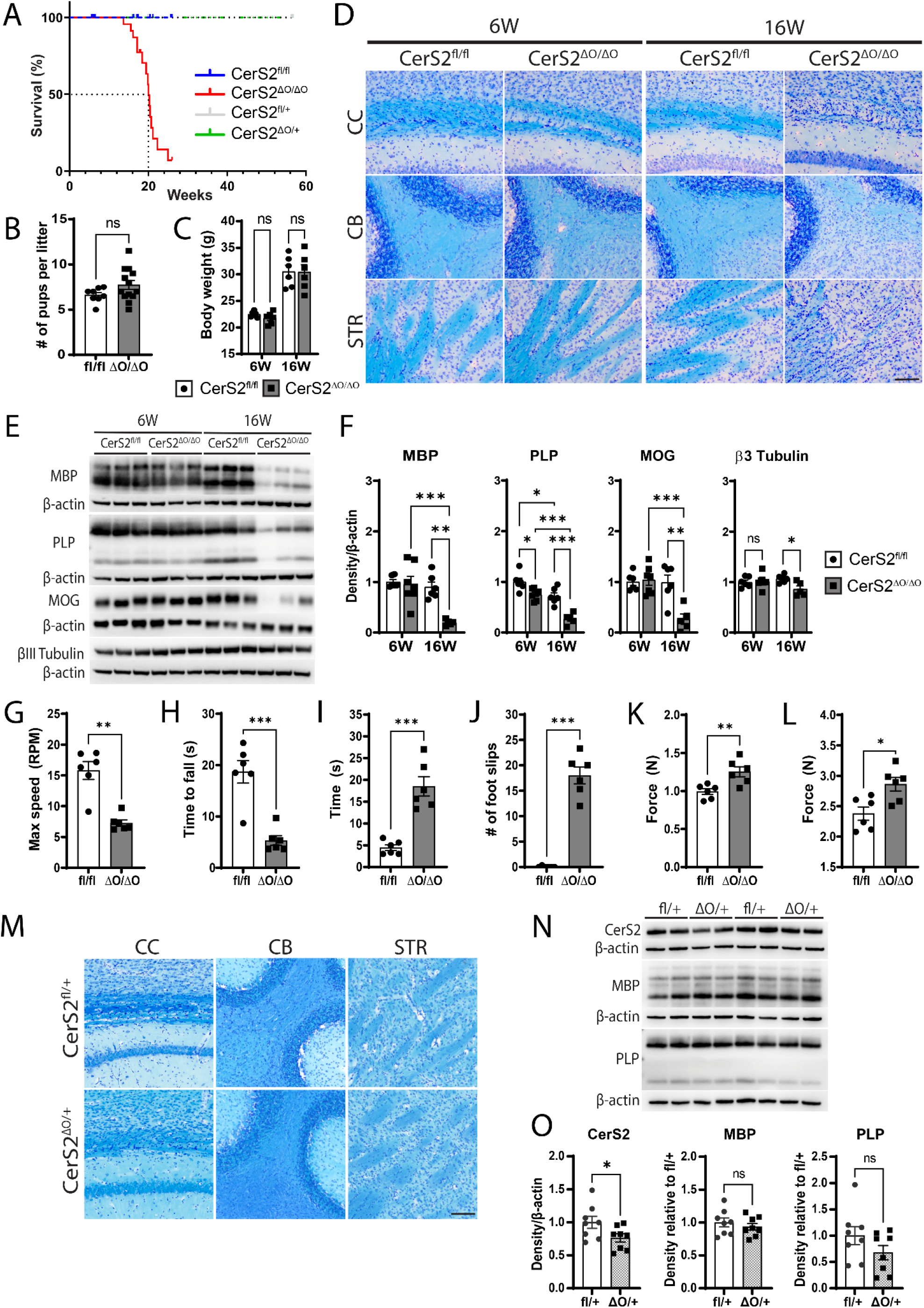
Myelin loss and impaired motor function in CerS2^ΔO/ΔO^ mice. (A) Survival curves for different mouse genotypes used in this study, including breeders (total 96 mice). (B) Average litter sizes of CerS2^fl/fl^ (n = 8) and CerS2^ΔO/ΔO^ mice (n = 13). (C) Mouse weight at 6 and 16 weeks (6W and 16W) of age (6 mice/group). Open bars represent CerS2^fl/fl^ mice and grey bars represent CerS2^ΔO/ΔO^ mice. (D) Representative luxol fast blue staining of corpus callosum (CC), cerebellum (CB) and striatum (STR) from 6W and 16W CerS2^fl/fl^ and CerS2^ΔO/ΔO^ mice. Scale bar: 100 µm. (E) Representative western blots and (F) densitometric quantification (6 mice/group) of MBP, PLP, MOG and β3 tubulin in half-brain homogenates. Protein levels were normalised to β-actin, then to the average of the 6W CerS2^fl/fl^ group. Data were analysed using two-way ANOVA (age and genotype as variables) with Bonferroni’s post-hoc test. (G) Maximum speed achieved and (H) average time on the rotarod prior to falling off, for 16W mice. (I) Average time taken to cross the balance beam and (J) number of foot slips on the beam for 16W mice. (K) Average grip strength for forelimbs alone or (L) both fore- and hindlimbs, 16W mice. Data in G-L are the average for 3 trials per mouse, presented as mean ± SEM. Statistical analyses were performed by unpaired t-test (6 mice/group). (M) Representative luxol fast blue staining in 16W CerS2^ΔO/+^ mice and CerS2^fl/+^ controls. (N) Representative western blots and (O) densitometric quantification (8 mice/group) for CerS2, MBP and PLP in CerS2^ΔO/+^ and CerS2^fl/+^ mice. Protein levels were normalised to β-actin, then to the average of the CerS2^fl/+^ group. Statistical analyses were performed by unpaired t-test; * *p*<0.05, ** *p*<0.01, *** *p*<0.001.

At 16 weeks, CerS2^ΔO/ΔO^ mice displayed significant motor deficits compared to controls. They performed significantly worse in the rotarod test (Fig. 4G-H), took longer to traverse the balance beam (Fig. 4I) and made more foot slips (Fig. 4J) than CerS2^fl/fl^ controls. Interestingly, the CerS2^ΔO/ΔO^ mice registered greater fore- and hindlimb grip strength compared to controls (Fig. 4K-L).

No discernible loss of myelin (Fig. 4M) or myelin protein markers (Fig. 4N-O) was observed in 16-week-old mice heterozygous for CerS2 deletion in oligodendrocytes (CerS2^ΔO/+^). These mice did not exhibit any deficits in the rotarod test or any change in grip strength, but did show a significant increase in time taken to cross the balance beam and number of foot slips on the beam (Supplementary Fig. 2A-F), indicative of a subtle motor deficit.

### Reduced myelin thickness and percentage of myelinated axons in CerS2^ΔO/ΔO^ mice

In agreement with the luxol fast blue staining and western blotting, electron microscopy showed normal-appearing myelin in the corpus callosum of 6W CerS2^ΔO/ΔO^ mice (Fig. 5A-B). However, quantitative analysis revealed a reduction in the percentage of myelinated axons (Fig. 5C). Smaller-calibre axons < 0.5 µm were particularly affected (Fig. 5D). The g-ratio [diameter of axon/(axon + myelin)] was increased across all axon diameters in CerS2^ΔO/ΔO^ mice (Fig. 5E-F), indicative of reduced myelin thickness. Mean axon diameter and the distribution of axons as a function of their diameter did not differ between CerS2^ΔO/ΔO^ and CerS2^fl/fl^ mice (Fig. 5G-H).

**Figure 5.**
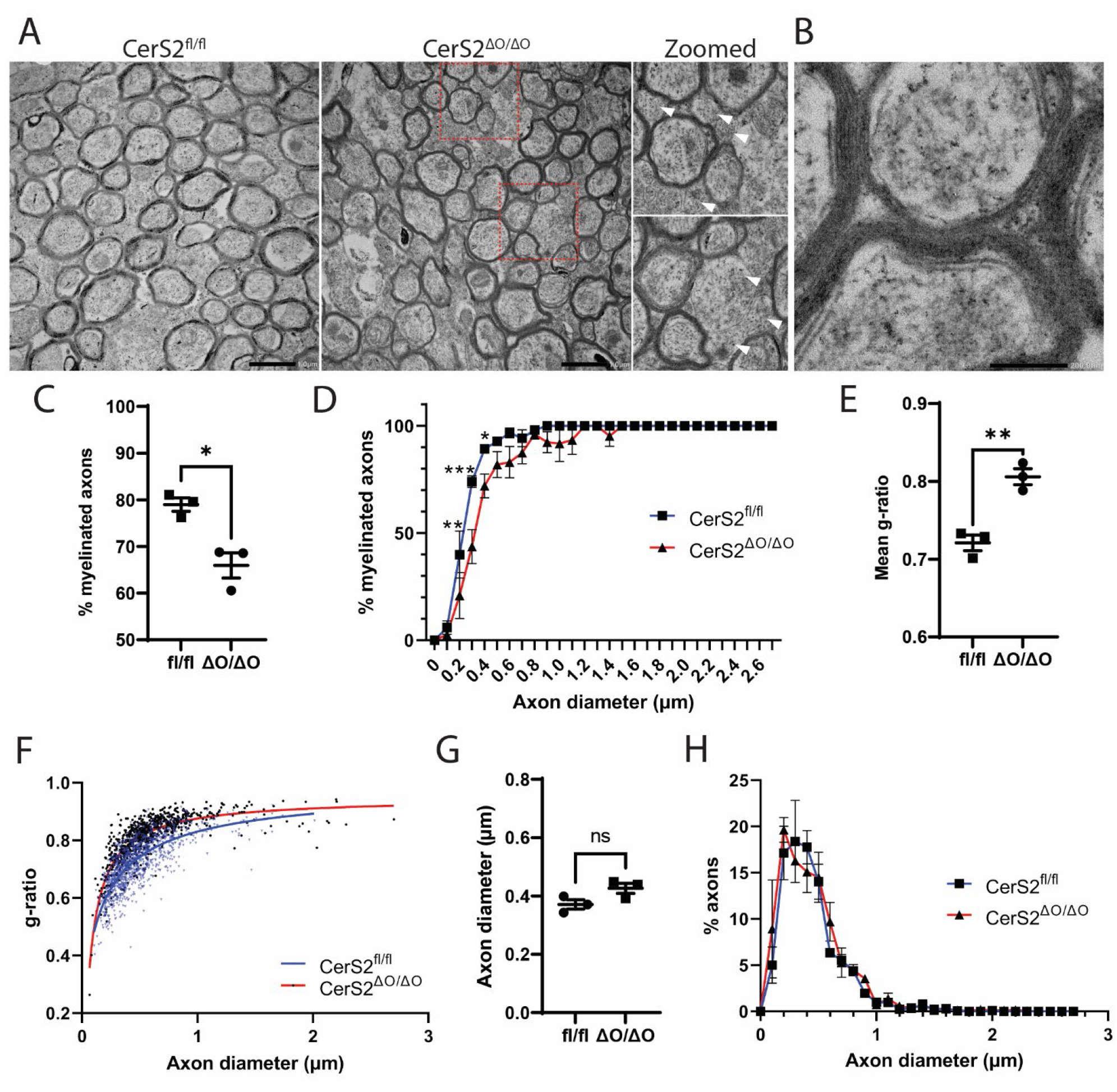
Myelin is thinner and absent from small-diameter axons in 6W CerS2^ΔO/ΔO^ mice. (A) Representative images of myelinated axons in CerS2^fl/fl^ and CerS2^ΔO/ΔO^ mice. Red bounding boxes correspond to the magnified images depicting unmyelinated axons (white arrow heads). Scale bar: 1 µm. (B) A magnified image of axons from a CerS2^ΔO/ΔO^ mouse depicting a thinner yet fully laminated axon adjacent to other thicker myelin sheaths. Scale bar: 200 nm. (C) Myelinated axons as a proportion of axons counted, (D) percentage of myelinated axons as a function of axon diameter, (E) mean g-ratio, and (F) g-ratio for each axon as a function of axon diameter. Curves of best-fit for CerS2^fl/fl^ (blue points and blue line) and CerS2^ΔO/ΔO^ mice (black points and red line) are shown. (G) Mean axon diameter and (H) percentage of axons falling within each axon diameter bin for CerS2^fl/fl^ (blue line) and CerS2^ΔO/ΔO^ (red line) mice. Data in (C, D, E, G, and H) is presented as mean ± SEM for each of 3 mice per group. Statistical analyses were performed by unpaired t-test in C, E, and G, and two-way ANOVA with Bonferroni’s post-hoc test in D and H; * *p*<0.05, ** *p*<0.01, *** *p*<0.001.

### Gliosis and axon stress precede oligodendrocyte loss in CerS2^ΔO/ΔO^ mice

We next sought to determine if loss of myelin in CerS2 mice is accompanied by loss of mature oligodendrocytes, by immunofluorescent labelling for the mature oligodendrocyte marker aspartoacylase (ASPA) (Madhavarao et al., 2004; Song et al., 2021; Zhang et al., 2014). At 6 weeks of age, there was no significant difference in the density of mature oligodendrocytes in CerS2^ΔO/ΔO^ compared to CerS2^fl/fl^ mice (Fig. 6A-B). By 16 weeks, mature oligodendrocyte density was 33% lower in the corpus callosum and 56% lower in the striatum of CerS2^ΔO/ΔO^ mice, although unchanged in the cerebellum.

**Figure 6.**
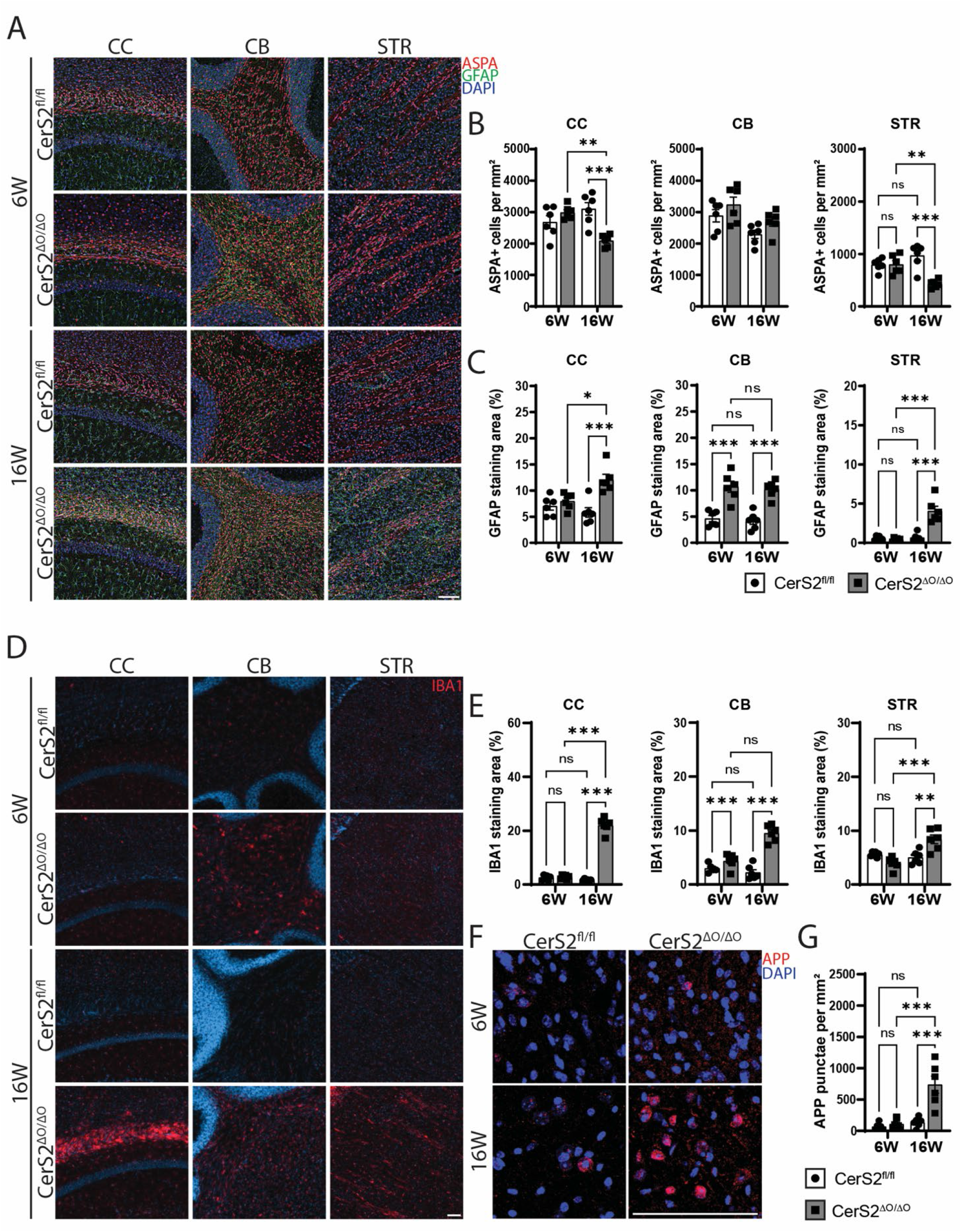
Oligodendrocyte loss, astrogliosis and microgliosis in 16W CerS2^ΔO/ΔO^ mice. (A) Representative images of mature oligodendrocytes (ASPA, red) and astrocytes (GFAP, green) in the corpus callosum (CC), cerebellum (CB) and striatum (STR) of 6W and 16W CerS2^fl/fl^ and CerS2^ΔO/ΔO^ mice. DAPI nuclear stain in blue. (B) Density of ASPA-positive cells in the CC, CB and STR of 6W and 16W CerS2^fl/fl^ and CerS2^ΔO/ΔO^ mice. (C) Area of staining (% of total area) for GFAP. (D) Representative images of IBA1 staining (microglia) in the CC, CB and STR of 6W and 16W CerS2^fl/fl^ and CerS2^ΔO/ΔO^ mice. (E) Area of IBA1 staining (% of total area) in the CC, CB and STR of 6W and 16W CerS2^fl/fl^ and CerS2^ΔO/ΔO^ mice. (F) Representative APP staining (red) with DAPI counterstain (blue), and (G) density of APP punctae in the CC. Open bars represent CerS2^fl/fl^ mice and grey bars represent CerS2^ΔO/ΔO^ mice (6 mice/group). Data are presented as mean ± SEM. Statistical analyses were performed by two-way ANOVA with Bonferroni’s post-hoc test. * *p*<0.05, ** *p*<0.01, *** *p*<0.001. Scale bar: 100 µm.

Myelin loss is often accompanied by proliferation and activation of astrocytes and microglia. These were measured by area of staining for Glial Fibrillary Acidic Protein (GFAP) and Ionized Calcium Binding Adapter Protein 1 (IBA1), respectively. Both GFAP (Fig. 6A,C) and IBA1 (Fig. 6D,E) staining were modestly but significantly increased in the cerebellum of CerS2^ΔO/ΔO^ relative to control mice at 6 weeks of age, whereas no difference was seen in the corpus callosum or striatum. At 16 weeks, pronounced GFAP and IBA1 staining was apparent in all three regions in CerS2^ΔO/ΔO^ mice. GFAP and IBA1 staining did not differ between CerS2^ΔO/+^ and CerS2^fl/+^ mice at 16 weeks of age (Supplementary Fig. 2G-I).

Axonal spheroids that stain intensely for amyloid precursor protein (APP) are an established indicator of axon injury (Hoflich et al., 2016; Sherriff et al., 1994) thought to mark sites of cytoskeletal breakdown (Sherriff et al., 1994). The extent of APP staining and number of APP-positive puncta were significantly increased in the corpus callosum of 16W CerS2^ΔO/ΔO^ mice (Fig. 6F-G).

Given that severe gliosis, myelin and oligodendrocyte loss were observed in 16- but not 6-week-old CerS2^ΔO/ΔO^ mice, we examined 10-week-old (10W) mice to narrow down the timeframe for these pathological changes. Myelin content by luxol fast blue staining did not show appreciable differences between CerS2^ΔO/ΔO^ and CerS2^fl/fl^ mice, although there were more cells in the white matter of CerS2^ΔO/ΔO^ mice (Fig. 7A). MBP levels were significantly lower in 10W CerS2^ΔO/ΔO^ mice, whereas PLP and MOG were not significantly different (Fig. 7B-C). Mature oligodendrocyte density did not differ between CerS2^ΔO/ΔO^ and CerS2^fl/fl^ mice (Fig. 7D-E), however GFAP and IBA1 staining were markedly increased in white matter of CerS2^ΔO/ΔO^ mice (Fig. 7F-I). APP puncta were clearly apparent in 10W CerS2^ΔO/ΔO^ mice (Fig. 7J-K). 10W CerS2^ΔO/ΔO^ mice did not exhibit motor deficits in the rotarod test and grip strength tests (Fig. 7L-O), but showed a delay in time taken to cross the balance beam (Fig. 7P) and an increased number of foot slips while traversing the beam (Fig. 7Q).

**Figure 7.**
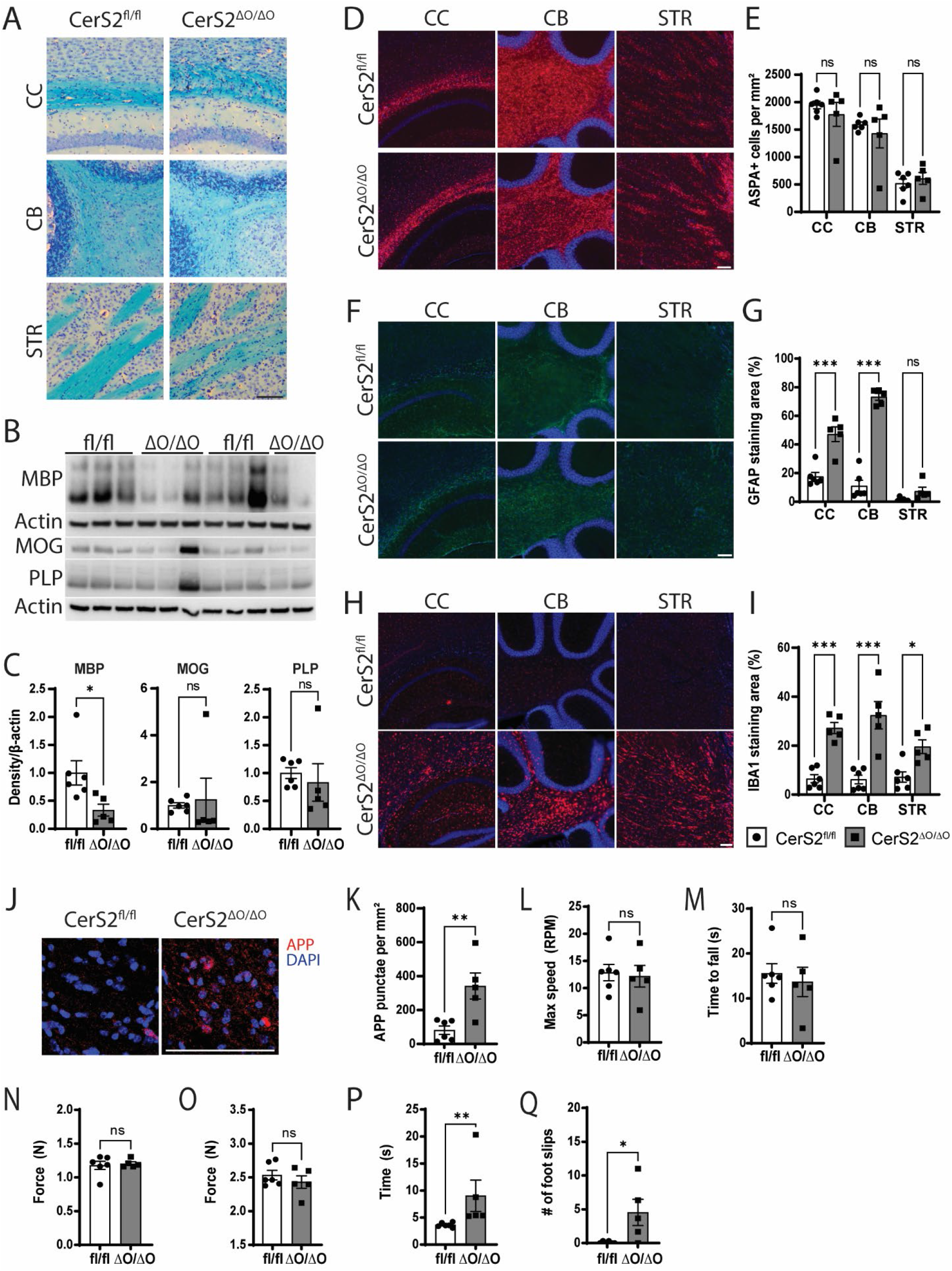
Astrogliosis and microgliosis without oligodendrocyte loss in 10W CerS2^ΔO/ΔO^ mice. (A) Representative luxol fast blue staining of corpus callosum (CC), cerebellum (CB) and striatum (STR) from 10-week-old (10W) CerS2^fl/fl^ and CerS2^ΔO/ΔO^ mice. (B) Representative western blots and (C) densitometric quantification of MBP, PLP, and MOG in half-brain homogenates. Myelin proteins were normalised to β-actin. (D) Representative staining for ASPA (red) and DAPI (blue), and (E) density of ASPA-positive cells in the CC, CB, and STR of 10W CerS2^fl/fl^ and CerS2^ΔO/ΔO^ mice. (F-I) Representative images and quantified area of staining for GFAP (F, G) and IBA1 (H, I). Statistical analyses were performed by two-way ANOVA with Bonferroni’s post-hoc test. (J) Representative APP staining with DAPI counterstain, and (K) density of APP puncta in the CC. (L) Maximum speed and (M) average time on the rotarod prior to falling off. (N) Average grip strength for forelimbs alone or (O) both fore- and hindlimbs. (P) Average time taken to cross, and (Q) number of foot slips on the balance beam. Data are presented as mean ± SEM. Open bars represent CerS2^fl/fl^ mice (n = 6), and grey bars represent CerS2^ΔO/ΔO^ mice (n = 5). Statistical analyses were performed by unpaired Student’s t-test or Mann-Whitney test, * *p*<0.05, ** *p*<0.01, *** *p*<0.001. Scale bar: 100 µm.

### A phagocytic microglial population is apparent at 6 weeks of age in CerS2^ΔO/ΔO^ mice

To determine whether microgliosis was associated with increased microglial cell number or a phenotypically-distinct population of cells, we used high-parameter spectral cytometry to quantify and phenotype immune cells in whole brains (Spiteri et al., 2021). The gating strategy is shown in Supplementary Fig. 3. Other than increased neutrophils in CerS2^ΔO/ΔO^ mice (F = 10.1, *p* = 0.013 for effect of genotype in 2-way ANOVA), there were no significant changes to immune cell numbers, including total microglia (Fig. 8A). Clustering and dimensionality reduction using FlowSOM and UMAP analysis on 12 phenotypic markers revealed three microglial clusters (Fig. 8B-C). These include the previously reported CD86- and CD86+ microglial subsets (Spiteri et al., 2022; Spiteri et al., 2021), as well as a distinct population that we have termed myelin-responsive microglia (MRM), prominent in CerS2^ΔO/ΔO^ but not CerS2^fl/fl^ mice (Fig. 8B-C). MRM exhibited downregulation of TMEM119 and P2RY12, which are generally thought of as homeostatic or resting microglial markers (Krasemann et al., 2017; Spiteri et al., 2021); and significant upregulation of CD11c and CD68, the latter being a lysosomal protein expressed by phagocytic microglia (Fig. 8D and Supplementary Fig. 4). The MRM population also expressed higher levels of other markers associated with immune responses, specifically CD45, MHC-II, CD11b, and CD64 (Fig. 8E) and was approximately 4-fold more abundant in 6-week-old CerS2^ΔO/ΔO^ compared to CerS2^fl/fl^ mice, increasing to 8-fold in 10-week-old CerS2^ΔO/ΔO^ mice (2-way ANOVA, effect of genotype: F = 39.6, *p* = 0.0002; age: F = 8.7, *p* = 0.019; interaction: F = 8.5, *p* = 0.020) (Fig. 8C).

**Figure 8.**
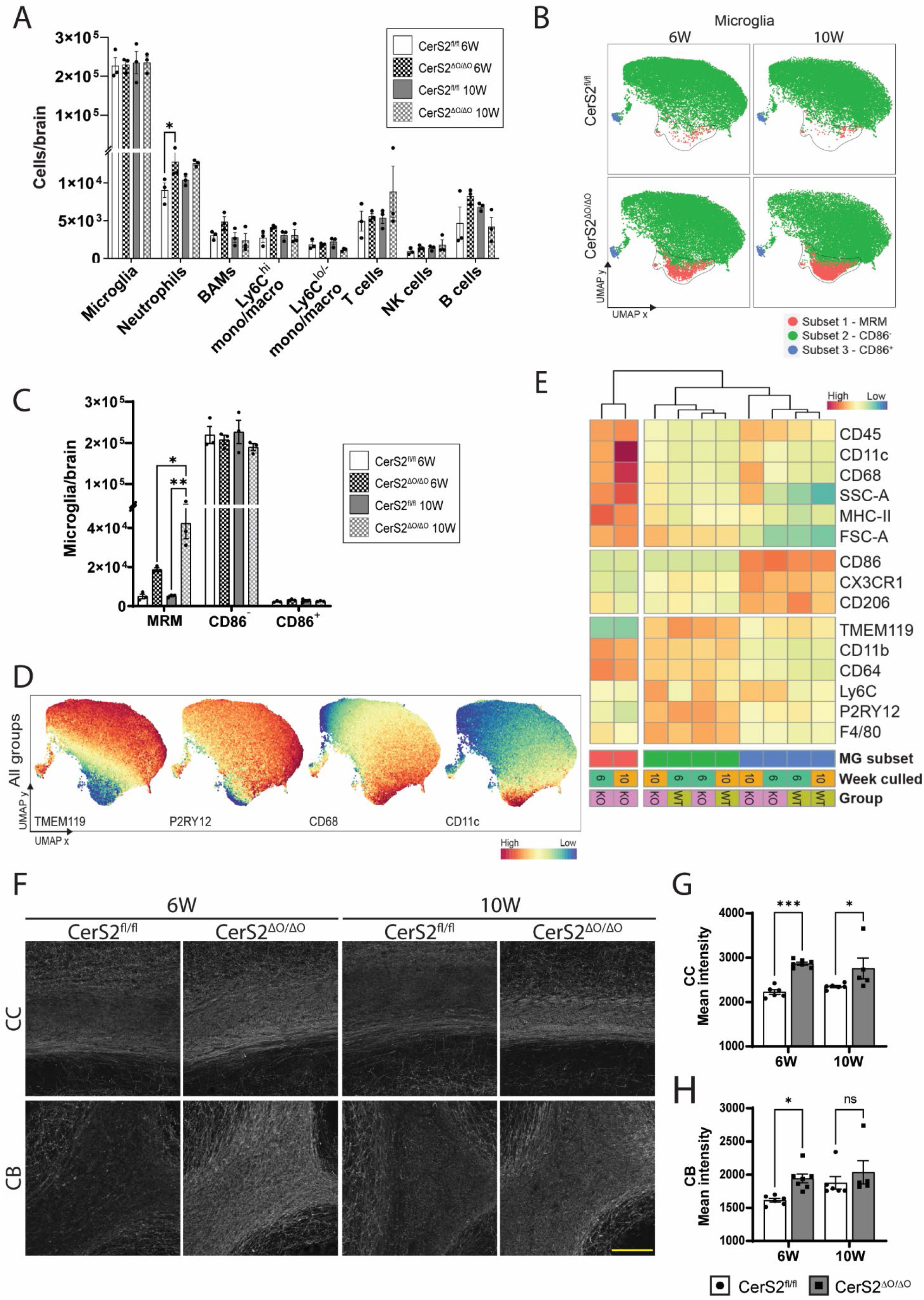
A distinct population of phagocytic microglia in CerS2^ΔO/ΔO^ mice. (A) Numbers of different immune cell types in whole brains of 6W and 10W CerS2^fl/fl^ and CerS2^ΔO/ΔO^ mice. (B) UMAP plots based on a panel of 12 markers expressed by microglia. A population highlighted in red and labelled myelin-responsive microglia (MRM) is increased in CerS2^ΔO/ΔO^ mice. (C) Number of microglia in the MRM, CD86^−^ or CD86^+^ populations shown in part (B). (D) UMAP plots showing the relative levels of TMEM119, P2RY12, CD68 and CD11c in the different microglial populations. (E) Heatmap showing mean expression level for 13 protein markers, as well as forward (FSC-A) and side scatter (SSC-A), in microglial populations shown in (B): MRM (red), CD86^−^ (green) and CD86^+^ (blue). Age at cull (6 weeks in teal, 10 weeks in orange) and genotype (CerS2^ΔO/ΔO^ in pink, CerS2^fl/fl^ in olive) are indicated below the heatmap. Data is presented as mean ± SEM (A and C), or mean (B, D, E) of 3 mice/group. (F) Representative staining in the corpus callosum (CC) and cerebellum (CB) of 6W and 10W mice. Scale bar: 100 µm. (G, H) Mean denatured MBP staining intensity in the CC (G) and CB (H) of CerS2^fl/fl^ (circles, 6 mice/group) and CerS2^ΔO/ΔO^ mice (squares, 6W: n=7, 10W: n=5). Statistical analyses in (A), (C), (G), and (H) were performed by two-way ANOVA (genotype and age as variables) followed by Bonferroni’s post-hoc test: * *p*<0.05, ** *p*<0.01 *** *p*<0.001.

Given the presence of this phagocytic microglial population, we performed immunostaining for denatured MBP using an antibody that recognises an epitope of MBP that is exposed in denatured myelin (Cantoni et al., 2015; Matsuo et al., 1997). Denatured MBP staining was significantly increased in the corpus callosum (2-way ANOVA, effect of genotype: F = 33.3, *p* < 0.0001; age: F = 0.007, *p* = 0.94; interaction: F = 2.3, *p* = 0.14) and cerebellum (2-way ANOVA, effect of genotype: F = 8.1, *p* = 0.010; age: F = 4.1, *p* = 0.057; interaction: F = 1.5, *p* = 0.24) of 6-week-old CerS2^ΔO/ΔO^ compared to CerS2^fl/fl^ mice (Fig 8F-H). At 10 weeks of age, denatured MBP staining was significantly higher in the corpus callosum but not cerebellum of CerS2^ΔO/ΔO^ mice.

## Discussion

In this study we establish that selective ablation of very long chain ceramide synthesis in oligodendrocytes produces a decrease in myelin sphingolipid acyl chain length, triggering a series of degenerative changes that result in death of the mice between 4 and 6 months of age. Loss of CerS2 in myelinating cells resulted in greatly reduced C22-C24 sphingolipids and increased C16-C18 sphingolipids in myelin. This was associated with an overall reduction in myelin sheath thickness, a decrease in the proportion of myelinated axons, and immunoreactivity for denatured MBP at 6 weeks of age. A distinct microglial population expressing high levels of CD68 and CD11c was also apparent at 6 weeks, together with significant microgliosis and astrogliosis in the cerebellum. By 10 weeks of age, microgliosis, astrogliosis, and APP spheroids were strongly increased in white matter, along with evidence of reduced myelin content and functional deficits in the balance beam. Pronounced demyelination, oligodendrocyte loss, and deficits in the rotarod became apparent at 16 weeks of age. Overall, our results demonstrate that sphingolipid acyl chain length is critical for myelin stability, as substitution of very long chain with long chain sphingolipids produced marked gliosis, phagocytic microglia that probably drive myelin degradation, and axon stress prior to young adulthood.

In the CNS, CerS2 is most highly expressed by mature oligodendrocytes (Becker et al., 2008; Couttas et al., 2016; Kremser et al., 2013; Zhang et al., 2014) and a prior study showed the CerS2 immunoreactivity was restricted to oligodendrocytes (Kremser et al., 2013). In agreement, we observed strong CerS2 immunoreactivity only in Olig2-positive cells, and staining was absent in CerS2^ΔO/ΔO^ mice. Western blotting showed near-complete loss of CerS2 protein in white matter, but only around 50% reduction in cortex, indicating that CerS2 is not exclusively expressed by oligodendrocytes in the CNS. However, levels of C22-C24 sphingolipids were substantially lower in whole brain homogenates of CerS2^ΔO/ΔO^ mice, confirming that oligodendrocytes are responsible for the bulk of very long chain ceramide synthesis in the brain.

In agreement with the ubiquitous CerS2 knockout (CerS2-null) model (Ben-David et al., 2011; Imgrund et al., 2009), myelin was formed in CerS2^ΔO/ΔO^ mice but deteriorated in the first four months of life. Prior studies with CerS2-null mice did not examine the structure or lipid composition of myelin in the weeks immediately following developmental myelination, which peaks at 3 weeks and is mostly complete by 6 weeks of age in mice (Semple et al., 2013). We show that fewer axons in the corpus callosum are myelinated in CerS2^ΔO/ΔO^ mice at 6 weeks of age, and myelin is thinner in those axons that are myelinated. Mature oligodendrocyte density was not affected at this age, indicating either that the inability to synthesize very long chain sphingolipids attenuates myelination by these oligodendrocytes, or that significant myelin degradation is already occurring. The latter is plausible given that a distinct population of phagocytic microglia were apparent at this age.

Prior studies with CerS2-null mice reported substantial loss of HexCer in white matter from 10 weeks of age (Ben-David et al., 2011; Imgrund et al., 2009), and suggested that encephalopathy is attributed to the reduction in HexCer content. In contrast, we observed a modest increase in total HexCer, ST, and ceramide in myelin of 6-week-old CerS2^ΔO/ΔO^ mice, attributed to increased C18 and, to a lesser extent, C16 sphingolipids that compensated quantitatively for loss of C22 and C24 variants. The difference is likely due to our examination of myelin at 6 weeks of age rather than white matter at 10 weeks or older, when overall loss of myelin will be reflected in reduced HexCer and ST content. Increased C16 and C18 sphingolipid content probably stems from increased dihydrosphingosine substrate availability for other CerS isoforms, particularly CerS1, following CerS2 deletion in oligodendrocytes. This increase in dihydrosphingosine is consistent with prior studies on CerS2-null mice (Pewzner-Jung, Park, et al., 2010). We confirmed that the C18 HexCer synthesized by CerS2^ΔO/ΔO^ mice is GalCer, not GluCer. This implies that the C18 HexCer is synthesized by oligodendrocytes, since other CNS cell types do not express ceramide galactosyltransferase (Schaeren-Wiemers et al., 1995; Zhang et al., 2014), and refutes the prior suggestion that ceramide galactosyltransferase cannot utilise C18 ceramide substrates (Ben-David et al., 2011). We therefore propose that redistribution of sphingolipid chains from C22/C24 to C16/C18 in newly formed myelin leads to myelin destabilisation. In support of this hypothesis, white matter of 6-week-old CerS2^ΔO/ΔO^ mice showed strong immunoreactivity for denatured MBP. This denatured MBP epitope is detected in demyelinating diseases and disease models, such as cuprizone feeding (Cantoni et al., 2015; Matsuo et al., 1997).

The very long acyl chains of myelin galactosphingolipids are probably critical for their self-segregation into raft-like, liquid-ordered domains, since this self-aggregation was lost with myelin lipids isolated from CerS2-null mice (Yurlova et al., 2011). This may disrupt the positioning of myelin proteins, since the association of PLP, MOG, and even the more soluble MBP with detergent-insoluble membrane fractions is dependent on GalCer and ST (Ozgen et al., 2016). Interdigitation of long C24 acyl chains in the outer leaflet with the inner leaflet membrane may also be important for creating raft-like myelin membrane domains (Pinto et al., 2008). Given that C22/C24 sphingolipids are replaced by C16/C18 variants in young CerS2^ΔO/ΔO^ mice, it would be valuable in future studies to determine whether their myelin lipids segregate into liquid-ordered domains and whether the association of myelin proteins with detergent-resistant lipid domains is preserved.

Recent research has established that lipid synthesis in astrocytes is essential for myelination and astrocyte lipid synthesis can compensate for a deficiency in oligodendrocyte lipid synthesis (due to CNP-Cre-mediated deletion of sterol regulatory element-binding protein (SREBP) cleavage-activating protein, SCAP) (Camargo et al., 2017). Deficiencies in cholesterol or fatty acid synthesis in oligodendrocytes delay myelination but do not cause myelin instability (Camargo et al., 2017; Saher et al., 2005). Our results indicate that this is not the case with very long chain sphingolipids. The essential requirement for CerS2 in oligodendrocytes may be attributed to the specifics of myelin sphingolipid synthesis: CerS2 is more highly expressed in oligodendrocytes than other CNS cells (Couttas et al., 2016; Kremser et al., 2013; Zhang et al., 2014), and only oligodendrocytes express the enzymes needed to synthesise GalCer and ST (Schaeren-Wiemers et al., 1995; Zhang et al., 2014). Mice unable to synthesize GalCer and ST, due to a deficiency in UDP-galactose ceramide galactosyltransferase, produce normal-appearing but thinner myelin that is functionally compromised, with sciatic nerve conduction speeds similar to those of unmyelinated axons. Similar to CerS2^ΔO/ΔO^ or CerS2-null mice, this myelin degenerates rapidly and the mice die young (Bosio et al., 1996; Coetzee et al., 1996).

Early lethality in CerS2^ΔO/ΔO^ mice implies that myelin loss and the associated gliosis are the cause of early lethality in the CerS2-null (ubiquitous knockout) mice (Pewzner-Jung, Brenner, et al., 2010). The mean age at death in CerS2-null mice was also around 5 months (Pewzner-Jung, Brenner, et al., 2010), however very few CerS2^ΔO/ΔO^ mice survived to 6 months, contrasting with the protracted lifespan of many CerS2-null mice. The more aggressive lethality could result from strain differences, as our CerS2^ΔO/ΔO^ mice were on a C57BL/6 background whereas published studies on CerS2-null mice used a mixed background (Imgrund et al., 2009; Pewzner-Jung, Park, et al., 2010). Death in CerS2^ΔO/ΔO^ mice was sudden and unexpected. This necessitated modification of our ethics protocol such that CerS2^ΔO/ΔO^ mice were only kept up to four months of age, and we were unable to determine the exact cause of death. Imgrund et al suggested that difficulty initiating motor function in CerS2-null mice may be attributed to neurodegeneration, due to the expression of *CERS2* mRNA in both neurons and oligodendrocytes (Imgrund et al., 2009). However, CerS2^ΔO/ΔO^ mice exhibited deficits in traversing the balance beam and impaired performance on the accelerod, indicating that these problems with balance and coordination result from loss of CerS2 in myelinating cells. The increased force required to break CerS2^ΔO/ΔO^ mice’s grip is potentially a consequence of myelin attrition affecting muscle tone. In other neurological injuries such as stroke, difficulty in releasing a grip is attributed to delays in terminating grip flexor muscle activity (Seo et al., 2009). People with MS have also been reported to exert significantly larger peak grip forces than healthy controls (Iyengar et al., 2009).

Microgliosis and astrogliosis were observed as early as 6 weeks of age in the cerebellar white matter of CerS2^ΔO/ΔO^ mice, indicating that these cells sensed and reacted to the altered myelin composition soon after developmental myelination. Myelination of the cerebellum commences early in development, which may explain why gliosis was evident earlier in this region (Buyanova & Arsalidou, 2021; Semple et al., 2013). More detailed phenotypic analysis identified a distinct population of microglia (MRM) in CerS2^ΔO/ΔO^ mice, expressing high levels of CD45, MHC-II, CD68, and CD11c, and reduced levels of Tmem119, Cx3CR1, and P2ry12 (Butovsky et al., 2014; Krasemann et al., 2017). MRM resemble microglia seen in mouse models of multiple sclerosis and Alzheimer’s disease (Ajami et al., 2018; Keren-Shaul et al., 2017; Krasemann et al., 2017). CD11c is highly expressed on microglia in post-natal development and neurodegenerative disease contexts (Benmamar-Badel et al., 2020). These CD11c-high microglia have been associated with a neuroprotective role, since deletion of CD11c-high microglia impaired myelination in the developing brain (Wlodarczyk et al., 2017), whereas their expansion reduced disease progression, demyelination and loss of oligodendrocytes in a mouse model of multiple sclerosis (Wlodarczyk et al., 2018). Up-regulation of the endo-lysosomal protein CD68 is a hallmark of phagocytic microglia (Cignarella et al., 2020; Schwabenland et al., 2021), suggesting that the MRM population expands in CerS2^ΔO/ΔO^ mice to facilitate clearance of myelin debris, which is required for remyelination (Cignarella et al., 2020; Zabala et al., 2018). Microglia are believed to mediate the bulk of homeostatic and pathological myelin phagocytosis in the CNS (Gomez-Sanchez et al., 2015), and myelin-laden microglia and macrophages are a feature of white matter diseases such as MS (Schwabenland et al., 2021). As such, understanding the role of phagocytic microglia may ultimately reveal important therapeutic opportunities.

Myelin phagocytosis by microglia can be initiated by interaction of the receptor Trem2 with myelin lipids such as ST and SM, when myelin or cellular debris is present (Cignarella et al., 2020; Poliani et al., 2015; Wang et al., 2015). This results in up-regulation of phagocytic markers and proteins that are traditionally associated with inflammatory activation, such as MHC-II and NOS2 (Cantoni et al., 2015), and downregulation of homeostatic markers (Krasemann et al., 2017). Trem2 sensing of myelin debris or lipids released from insecure myelin sheaths is a potential mechanism through which sphingolipid imbalance in CerS2^ΔO/ΔO^ mice triggers rapid demyelination. Resident glial cells may also sense abnormal myelin protein epitopes such as the denatured MBP epitope to which we demonstrated significantly-increased immunoreactivity. Alternatively, oligodendrocytes or neurons could emit a stress signal that initiates microglial and astrocyte activation. The tolerogenic function of microglia in suppressing T-cell activation and proliferation (Bai et al., 2009) is lost in activated microglia from neurodegenerative disease models (Krasemann et al., 2017). We observed no significant lymphocyte infiltration in the brains of 6W or 10W CerS2^ΔO/ΔO^ mice that might be suggestive of autoimmunity. In contrast, neutrophils were increased, as occurs in stroke and multiple sclerosis (Naegele et al., 2012; Perez-de-Puig et al., 2015).

Overall, our results support prior studies indicating that perturbations to myelin sphingolipid content adversely affect myelin stability (Ben-David et al., 2011; Bosio et al., 1996; Coetzee et al., 1996; Imgrund et al., 2009; Marcus et al., 2006). We extend these studies by demonstrating that a reduction in sphingolipid acyl chain length from C24 to C18, rather than loss of GalCer and ST, is sufficient to destabilise myelin and initiate the expansion of a distinct microglial population that likely phagocytose the myelin, leading to pronounced hypomyelination by 4 months of age. Our work also establishes that loss of very long chain sphingolipid synthesis in myelinating cells is of itself sufficient to cause neurological deficits, axon pathology, and early death, and cannot be compensated by synthesis of these lipids in other neural cells. Defining the triggers and receptors involved in microglial sensing of compromised myelin is tremendously important for understanding the pathogenesis of multiple sclerosis and other demyelinating diseases, and to inform new interventions that alleviate myelin degeneration.

## Acknowledgements

This research was supported by project grant APP1100626 and Ideas grant APP2002660 (A.S.D.) from the National Health and Medical Research Council (NHMRC), Australia; project grant 20-0113 from MS Research Australia (A.S.D. and L.P.); and an Australian government Research Training Stipend (O.C.M.). We gratefully acknowledge subsidised access to the Sydney Mass Spectrometry and Sydney Microscopy and Microanalysis core facilities, and subsidised mouse housing provided by Laboratory Animal Services, University of Sydney. Transmission electron microscopy sample preparation and imaging was performed at the Centre for Advanced Histology and Microscopy, Peter MacCallum Cancer Centre, Melbourne. The corresponding author is very grateful to Dr Yael Pewzner-Jung and Prof Anthony Futerman from the Weizmann Institute, Israel, for providing tissue samples from CerS2 knockout mouse that triggered the initiation of this project.

---

**Supplementary Figure 1.**
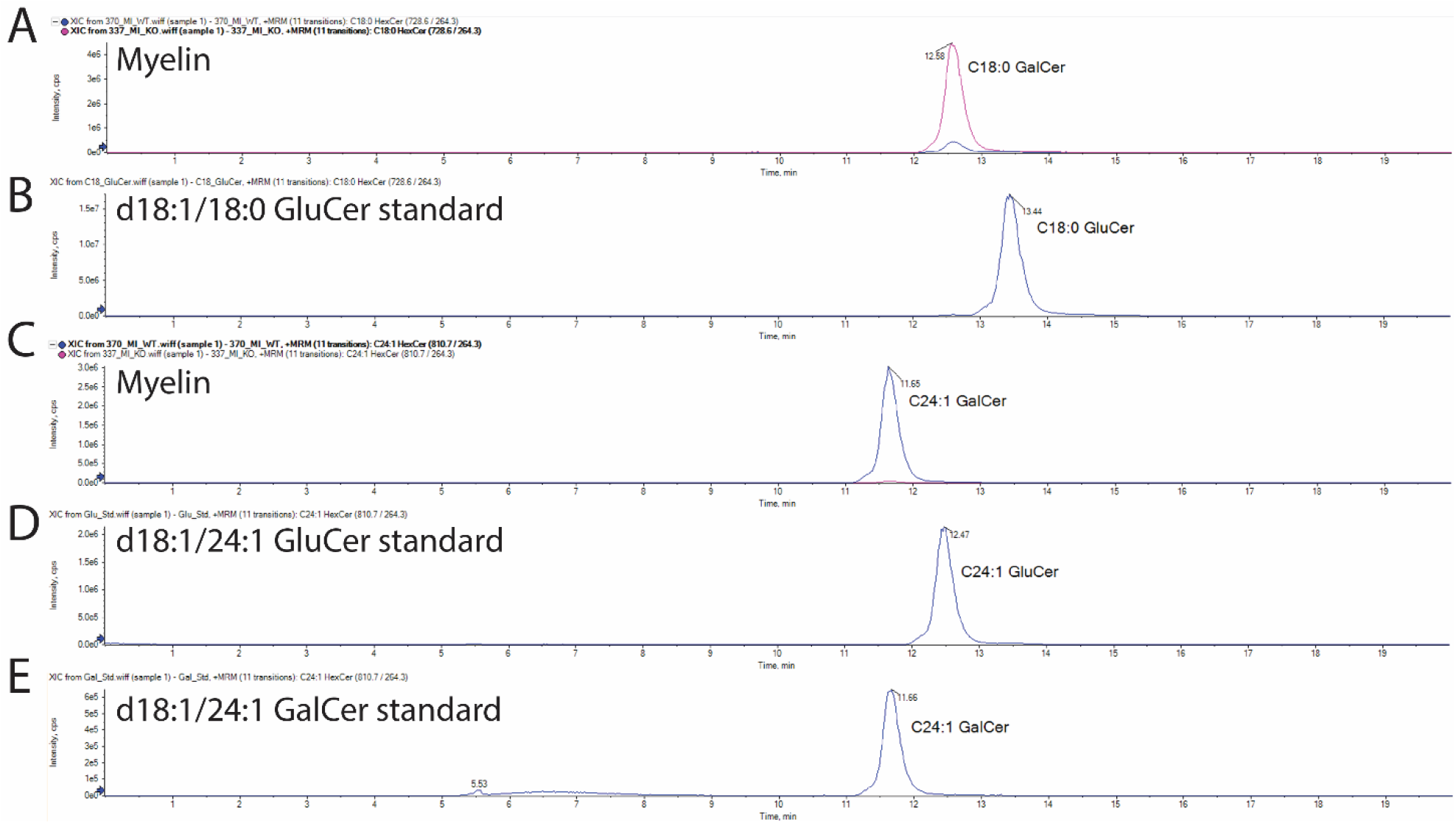
Myelin HexCer content is comprised wholly of GalCer. Chromatograms show (A,B) HexCer (d18:1/18:0) and (C-E) HexCer (d18:1/24:1) peaks in purified myelin from a CerS2^ΔO/ΔO^ (pink) and a CerS2^fl/fl^ (blue) mouse (A, C), a synthetic GluCer (d18:1/18:0) standard (B), a synthetic GluCer (d18:1/24:1) standard (D), and a synthetic GalCer (d18:1/24:1) standard (E). Peaks corresponding to GalCer but not GluCer were observed in all myelin samples. The GalCer (d18:1/18:0) peak was higher and GalCer (d18:1/24:1) much lower in myelin from CerS2^ΔO/ΔO^ mice, compared to CerS2^fl/fl^.

**Supplementary Figure 2.**
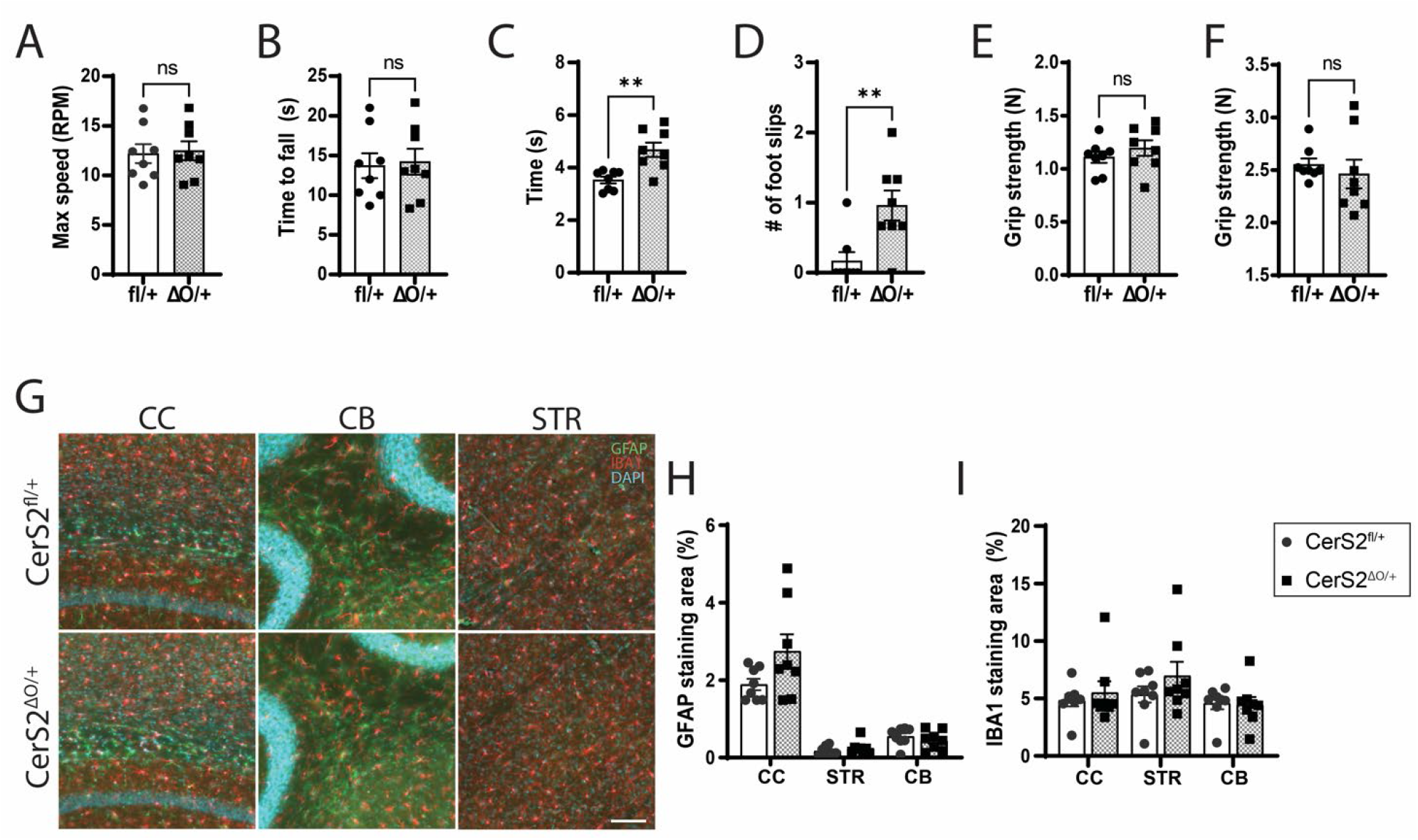
Motor function tests and gliosis in 16-week-old CerS2^ΔO/+^ mice. (A) Speed at which mice fell off the rotarod, and (B) time taken to fall off. (C) Time taken to cross the balance beam, and (D) the average number of foot slips made when crossing the beam. (E) Forelimb, and (F) forelimb + hindlimb grip strength. Results in A-F are the mean of 3 trials per mouse. (G) Representative images and area of staining for (H) GFAP (green) and (I) IBA1 (red) in the corpus callosum (CC), cerebellum (CB), and striatum (STR). DAPI nuclear stain in blue. Data are presented as mean ± SEM (8 mice per group). Statistical analyses were performed by unpaired t-tests or Mann-Whitney tests, ** *p*<0.01.

**Supplementary Figure 3.**
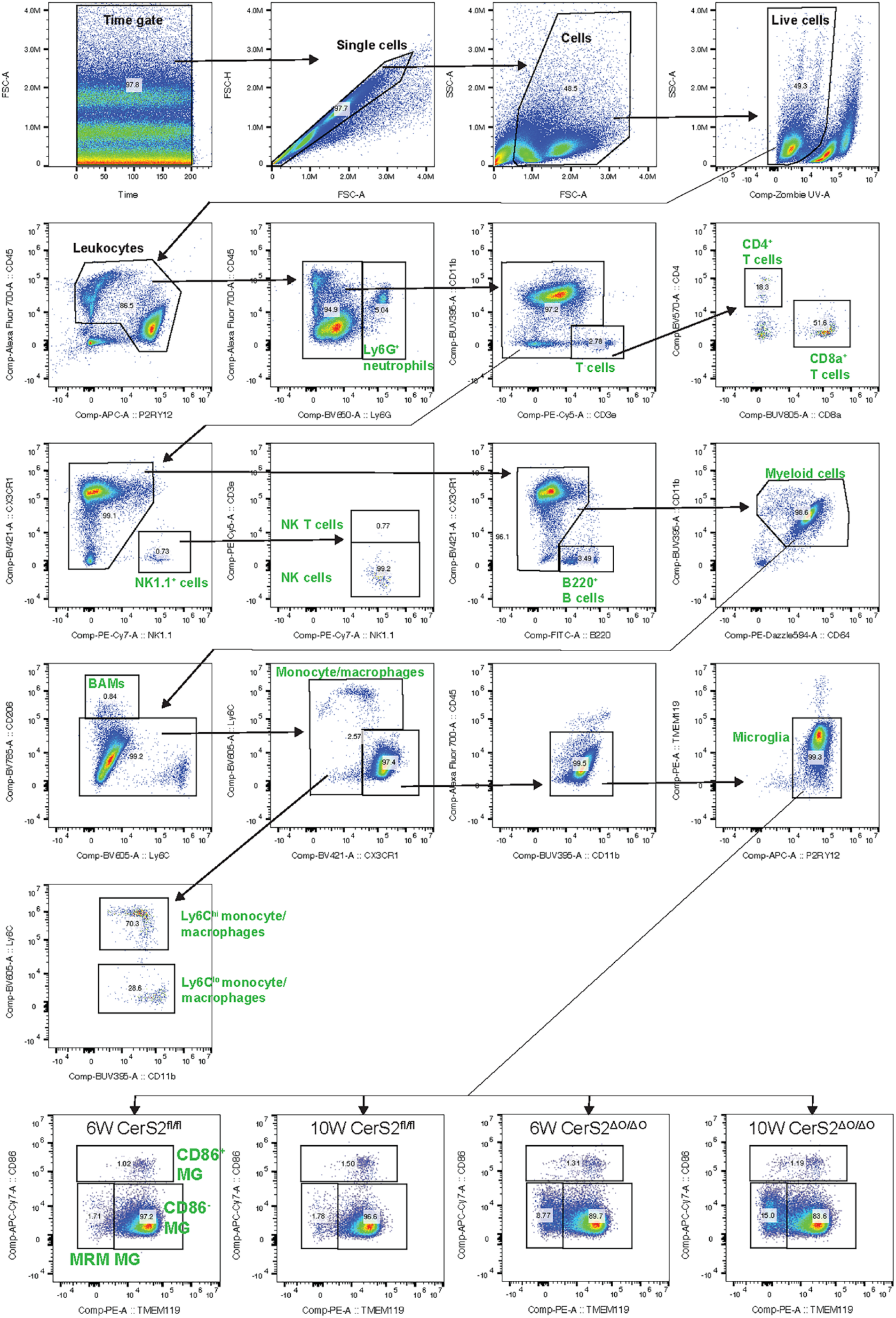
Flow cytometry gating scheme. Cell populations defined by specific markers are labelled in green font. MG: microglia.

**Supplementary Figure 4.**
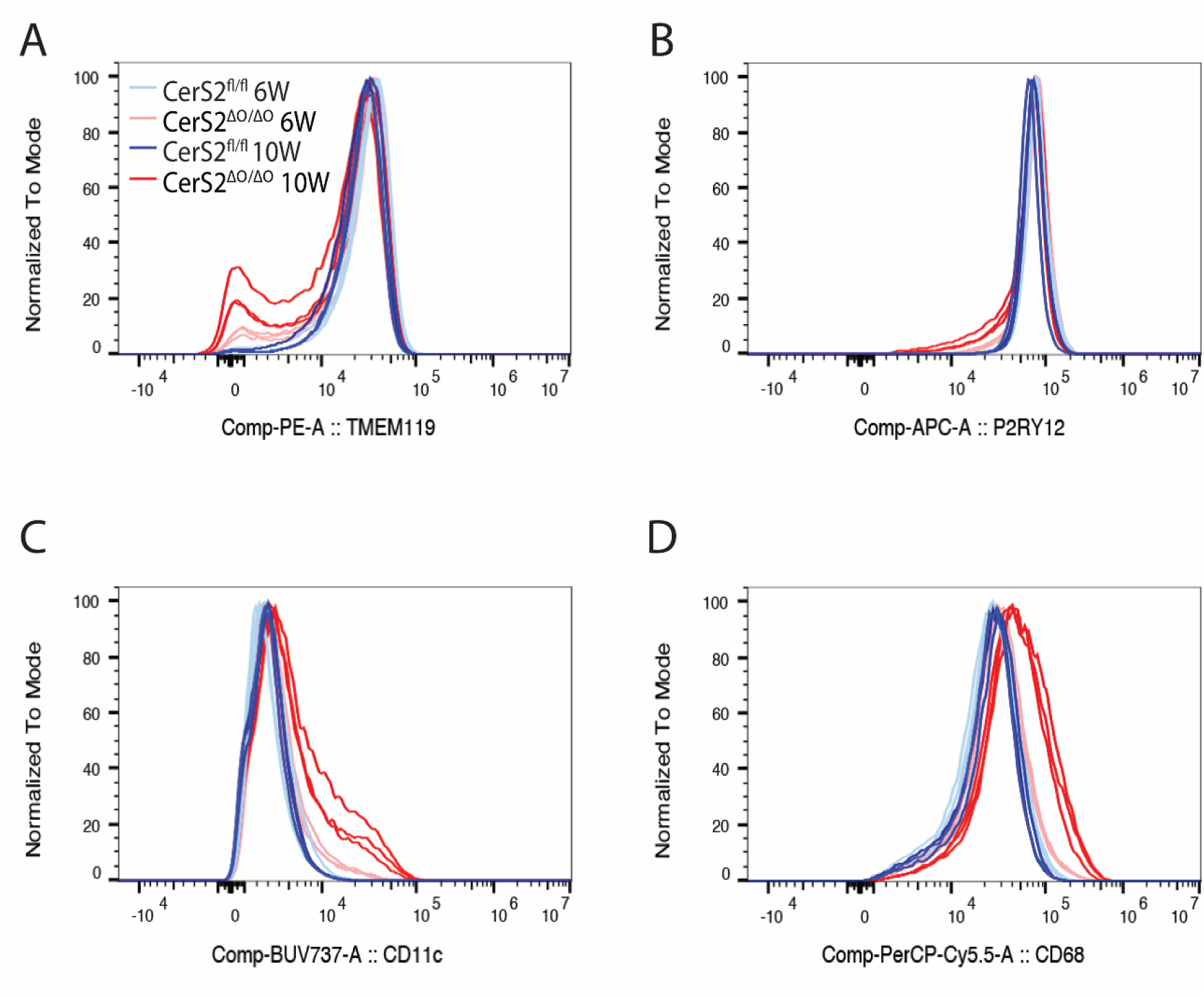
Expression markers for microglia. Histograms showing normalised cell number as a function of fluorescence intensity for (A) TMEM119 (B) P2RY12 (C) CD11c and (D) CD68 in microglia of 6-week-old CerS2^fl/fl^ (light blue), 10-week-old CerS2^fl/fl^ (dark blue), 6-week-old CerS2^ΔO/ΔO^ (pink), and 10-week-old CerS2^ΔO/ΔO^ (red) mice (3 mice/group).

